# SOX15 regulates stem cell pluripotency and promotes neural fate during differentiation by activating *Hes5*

**DOI:** 10.1101/2022.04.25.489402

**Authors:** Eun-Bee Choi, Munender Vodnala, Prince Saini, Madeleine Zerbato, Jaclyn J. Ho, Sharath Anugula, Jianing Wang, Shannan J. Ho Sui, Joon Yoon, Carla Inouye, Yick W. Fong

## Abstract

SOX2 and SOX15 are Sox family transcription factors enriched in embryonic stem cells (ESCs). The role of SOX2 in activating gene expression programs essential for stem cell self-renewal and acquisition of pluripotency during somatic cell reprogramming is well-documented. However, the contribution of SOX15 to these processes is unclear and often presumed redundant with SOX2 largely because overexpression of SOX15 can partially restore self-renewal in SOX2-deficient ESCs. Here, we show that SOX15 contributes to stem cell maintenance by cooperating with ESC-enriched transcriptional coactivators to ensure optimal expression of pluripotency-associated genes. We demonstrate that SOX15 depletion compromises reprogramming of fibroblasts to pluripotency which cannot be compensated by SOX2. Ectopic expression of SOX15 promotes the reversion of a post-implantation, epiblast stem cell state back to a pre-implantation, ESC-like identity even though SOX2 is expressed in both cell states. We also uncover a role of SOX15 in lineage specification, by showing that loss of SOX15 leads to defects in commitment of ESCs to neural fates. SOX15 promotes neural differentiation by binding to and activating a previously uncharacterized distal enhancer of a key neurogenic regulator, *Hes5*. Together, these findings identify a multifaceted role of SOX15 in induction and maintenance of pluripotency and neural differentiation.

## Introduction

Sox family transcription factors (TFs) play essential roles in development (1, 2). Sox TFs are characterized by their high-mobility group (HMG) DNA-binding domain that recognizes highly conserved DNA sequences (3). Despite having similar sequence specificity, Sox TFs are able to choreograph complex and divergent developmental processes by driving specific gene expression programs. In addition to their tissue-specific expression patterns, Sox proteins regulate cell type-specific transcription by leveraging their ability to partner with different TFs and recruit cell-specific transcriptional coactivators (4–6).

In embryonic stem cells (ESCs), SOX2 along with TFs OCT4 and NANOG activates a core regulatory network that is essential for stem cell self-renewal and pluripotency (7–11). SOX2 activates transcription by recruiting OCT4 and stem cell-enriched coactivators to gene enhancers containing sox-oct sequence motifs (12–15). The central role of SOX2 in reprogramming somatic cells to induced pluripotent stem cells (iPSCs) further highlights the importance of SOX2 in pluripotency (16–19). It has been shown that defects in stem cell self-renewal and iPSC generation in the absence of SOX2 can be rescued by ectopic expression of SOX1, SOX3, and SOX15 with varying degree of efficiency (20–22). However, the physiological relevance of the apparent functional redundancy is unclear, given that these SOX proteins were overexpressed in the rescue experiments, and that, except for SOX15, they are not naturally expressed in ESCs (23). In vitro binding assays indicate that SOX2 and SOX15 display essentially identical binding sequence preferences (23). Furthermore, OCT4 can pair with SOX2 or SOX15 and assemble onto an oct-sox enhancer element in vitro with comparable affinity (24). Another study showed by chromatin immunoprecipitation (ChIP) that several SOX proteins including SOX15 occupy oct-sox enhancers at select pluripotency genes that are also targeted by SOX2 (25). However, it is not known whether SOX15’s interaction with endogenous enhancers is functionally important for transcriptional activation. Therefore, the mechanism by which SOX15 contributes to the pluripotency gene network remains unclear.

When ESCs exit from pluripotency and undergo differentiation, collapse of the pluripotency gene network is accompanied by activation of lineage-specific differentiation programs. Pluripotency-associated TFs have been shown to play an active role in orchestrating this critical transcriptional switch (26). For example, SOX2 is redirected to neurogenic genes to promote neural fate specification (27, 28). The role of SOX15 in lineage specification is unknown.

In this study, we find that SOX2 depletion in ESCs leads to loss of SOX15 expression, raising important questions about whether defects in pluripotency gene expression and self-renewal found in SOX2-deficient ESCs could be due to simultaneous loss of SOX15. We therefore address the function of SOX15 in stem cell self-renewal, induction of pluripotency, and lineage specification. Using complementary in vitro biochemical approaches and genome-wide analyses, we show that SOX15 can partner with OCT4 to directly activate transcription of key pluripotency genes such as *NANOG*, by recruiting ESC-enriched transcriptional coactivators that are also targeted by SOX2. We show that ESCs depleted of SOX15 display defects in self-renewal and expression of pluripotency-associated genes despite having normal levels of SOX2. Likewise, we find that SOX15 depletion blocks reacquisition of pluripotency during cellular reprogramming which cannot be compensated by SOX2. Finally, we demonstrate that in differentiating ESCs, SOX15 is required for neural fate specification via direct binding and activation of a previously uncharacterized enhancer upstream of a key neurogenic gene, *Hes5*. Our findings identify common mechanisms of transactivation by SOX15 and SOX2 and reveal new non-redundant functions of SOX15 in induction of pluripotency and neural differentiation.

## Results

### OCT4 can cooperate with SOX2 or SOX15 to directly activate *NANOG* transcription

To address whether SOX15 and other related SOX proteins can cooperate with OCT4 to activate transcription, we took advantage of our fully reconstituted in vitro transcription assay that recapitulates stem cell-specific transcriptional activation by OCT4 and SOX2 (12). We expressed and purified a panel of SOX proteins (SOX1, SOX2, SOX11, and SOX15) and asked whether they can activate a well-characterized OCT4/SOX2-target, the human *NANOG* gene promoter (Figure 1 and figure S1A) (29, 30). In the presence of a partially purified pluripotent cell nuclear fraction (phosphocellulose 1M [P1M]) previously shown to contain transcriptional coactivators for OCT4/SOX2 (12), we found that, in addition to SOX2, SOX15 was able to robustly activate the *NANOG* gene with OCT4 (Figure 1A). Although it has been shown that SOX11 binds the enhancer of the *Nanog* gene in mouse ESCs (25), our result indicates that it is likely inactive for transcriptional activation, thus highlighting the selective nature of SOX-OCT4 protein complexes for stem cell-specific transcriptional activation. To confirm that SOX15 indeed binds sox-oct enhancers of the endogenous *Nanog* promoter and core pluripotency genes *Oct4* and *Sox2*, we used CRISPR/Cas9-mediated homologous recombination to knock-in an HA-epitope tag at the C-terminus of *Sox15* or *Sox2* gene locus in mouse ESCs (Figure 1B and Figure S1B). Binding of SOX15 at the sox-oct enhancers at *Oct4, Sox2*, and *Nanog* was confirmed by performing micrococcal nuclease (MNase)-chromatin immunoprecipitation (ChIP) using a highly specific anti-HA antibody (Figure 1C and figure S1C).

**Figure 1.**
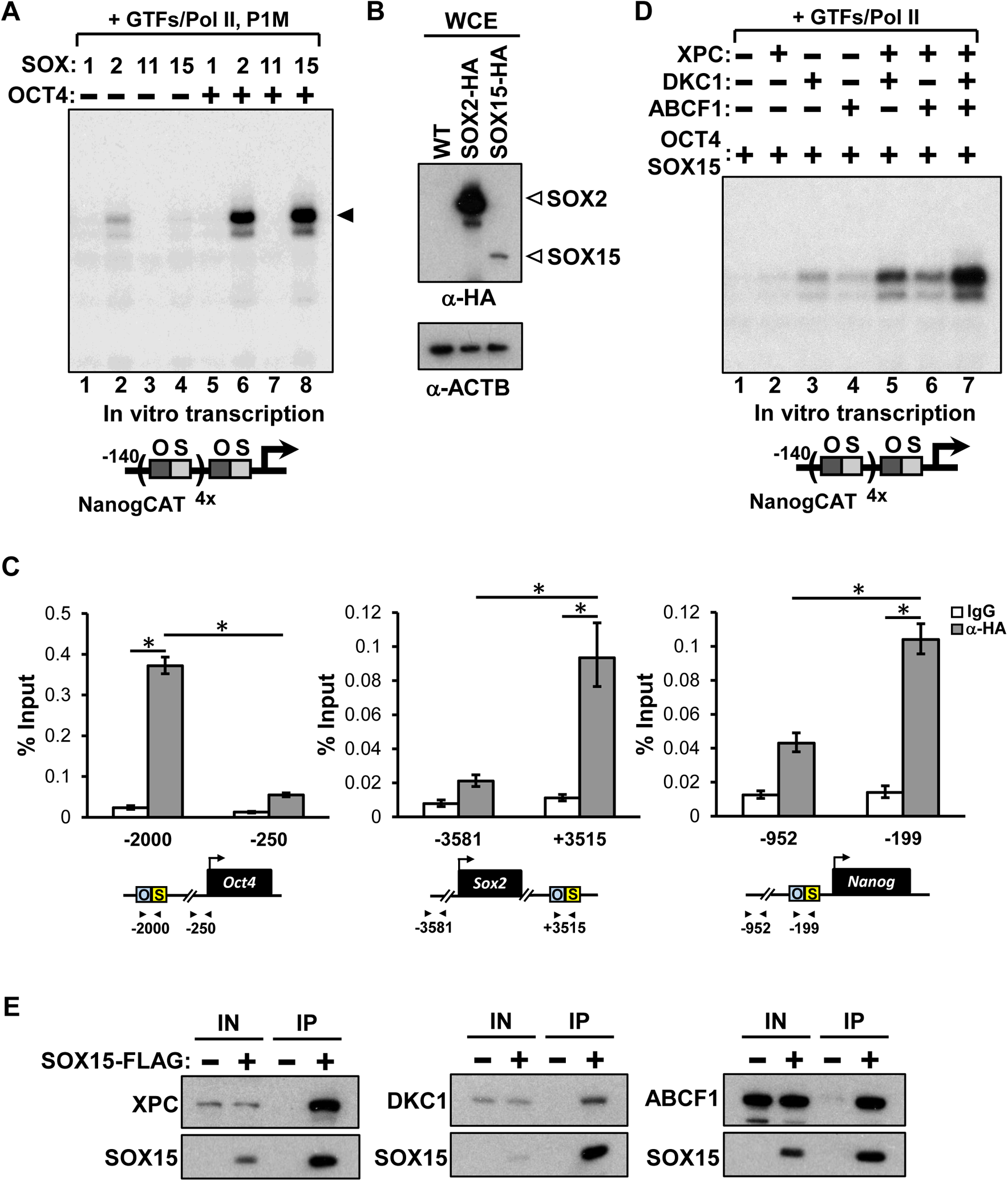
Transcriptional activation of *NANOG* by SOX2 and SOX15 requires Stem Cell Coactivators (SCCs). **(A)** Equal molar of indicated SOX proteins are added to in vitro transcription reactions in the absence (−) or presence (+)of OCT4 and tested for their ability to stimulate transcription from the human *NANOG* promoter template engineered with four extra copies of the oct-sox enhancer element (bottom). All reactions contain purified general transcription factors (GTFs), RNA polymerase II (Pol II), and a partially purified nuclear fraction (phosphocellulose 1M, [P1M]) containing stem cell-enriched coactivators (SCCs) essential for SOX2/OCT4-dependent transcriptional activation. Transcribed RNA products are subjected to primer extension and visualized by autoradiography. Filled arrowhead indicates major transcription products. **(B)** Expression of SOX2-HA and SOX15-HA in whole-cell extracts (WCEs) of knockin mouse ESC lines is analyzed by western blotting using an antibody against HA tag. β-actin (ACTB) is used as loading control. **(C)** Micrococcal nuclease (MNase) ChiP analysis of SOX15 occupancy on control and enhancer regions of *Oct4, Sox2,* and *Nanog* gene loci in SOX15-HA knockin mouse ESCs. Representative data showing the enrichment of SOX15-HA (gray bars) compared to control lgGs (white bars) are analyzed by qPCR and expressed as percentage of input chromatin. Schematic diagrams of known oct-sox binding sites of each gene and the relative positions of the amplicons are shown at the bottom. Error bars represent SEM. *n* = 3. (*) *P* < 0.05, calculated by two-sided Student’s t-test. (**D)** Transcriptional activation of *NANOG* by SOX15 and OCT4 in the presence of purified recombinant SCCs (XPC, DKC1, and ABCF1). **(E)** WCEs from 293T cells cotransfected with plasmids expressing FLAG-tagged SOX15 together with XPC, DKC1, or ABCF1 are immunoprecipitated with anti-FLAG antibody. Proteins coimmunoprecipitated with SOX15 are detected by western blotting using antibodies against XPC, DKC1, FLAG tag for SOX15-FLAG, and V5 tag for ABCF1-V5.

We have previously shown that transcriptional activation by SOX2 and OCT4 requires three stem cell coactivators (SCCs) present in P1M fraction, namely, the XPC DNA repair complex, the dyskerin (DKC1) ribonucleoprotein complex, and the ATP-binding cassette subfamily F member 1 (ABCF1) (12, 13, 15). We therefore asked whether SOX15 also potentiates transcription by cooperating with SCCs. We found that optimal transcriptional activation of *NANOG* requires all three SCCs (Figure 1D and figure S1D). The co-dependent nature of SCCs on transcriptional activation by SOX15 is highly reminiscent of our previous findings with SOX2 (13, 15), suggesting that SOX2 and SOX15 employ similar mechanisms to potentiate *NANOG* transcription in ESCs. We were unable to reproducibly detect interaction between SOX15 and SCCs in SOX15-HA knockin mouse ESC extracts by coimmunoprecipitation. We surmise that it could be due to the low expression level of SOX15 (see Figure 1B) and the often weak and transient nature of functional coactivator-activator interactions (31) Therefore, we tested SOX15-SCCs interactions by performing transient transfection and coimmunoprecipitation in 293T cells. We found that SOX15, like SOX2, binds XPC, DKC1, and ABCF1 (Figure 1E) (13, 15, 32, 33). Together with our previous observations that SCCs are recruited to SOX15-target genes including *Nanog*, *Oct4*, and *Sox2* in mouse ESCs (12, 13, 15), we propose that SOX15 activates stem cell-specific transcription by cooperating with SCCs through direct recruitment.

### SOX15 contributes to stem cell self-renewal and maintenance

To examine the functional relationship between SOX2 and SOX15 in ESCs, we performed acute depletion of SOX2 or SOX15 in mouse ESCs by targeted small hairpin RNAs (shRNAs) and assessed their effect on pluripotency gene expression. While knockdown of SOX15 has no discernible effect on SOX2 expression, we found that partial knockdown of SOX2 reduced SOX15 both at the mRNA and protein level (Figure 2A and 2B). These observations also emphasize the importance of studying SOX15 function in ESCs directly, because defects in transcription and pluripotency previously shown in SOX2-deficient ESCs may be attributed to loss of SOX15 (25, 34, 35). Consistent with previous observations, both SOX2 and SOX15 are enriched in the pluripotent state (Figure S2A) (23). Even though SOX15 mRNA and protein levels are significantly lower than that of SOX2 (Figure 1B and figure S2A), we found that knockdown of SOX15 was nevertheless able to reduce the expression of several key pluripotency genes such as *Nanog* and *Klf4*, although to a lesser degree than SOX2-knockdown (Figure S2B). These data suggest that both SOX2 and SOX15 contribute to the expression of these genes in ESCs. To determine whether incomplete nature of shRNA-mediated knockdown might obscure the impact of SOX15 depletion in ESCs, we employed CRISPR/Cas9 to knockout *Sox15* in mouse ESCs. In one knockout ESC line (KO1), frameshift indels were introduced near the 5’ end of *Sox15* coding region, leading to a premature translation stop at amino acid 72 (Figure 2C). Because the *Sox15* gene is only ~1.4 kb long, we also deleted *Sox15* by using two sgRNAs flanking the entire gene locus (KO2). Western blot analysis confirmed the absence of SOX15 in both KO ESC lines while SOX2 and OCT4 remained largely unchanged (Figure 2D). The fact that we could establish SOX15-null ESC lines indicated that self-renewal can occur in the absence SOX15. However, we noticed that when these SOX15-deficient ESCs were challenged with single cell dissociation, they were less efficient in forming pluripotent, alkaline phosphatase (AP)-positive cell colonies (Figure 2E). Consistent with their reduced self-renewal capacity, we found that the expression levels of several key pluripotency genes such as *Nanog*, *Klf4*, *Prdm14,* and *Esrrb* are compromised (Figure 2F). Therefore, our data suggested that optimal expression of these pluripotency genes and stem cell maintenance require both SOX15 and SOX2.

**Figure 2.**
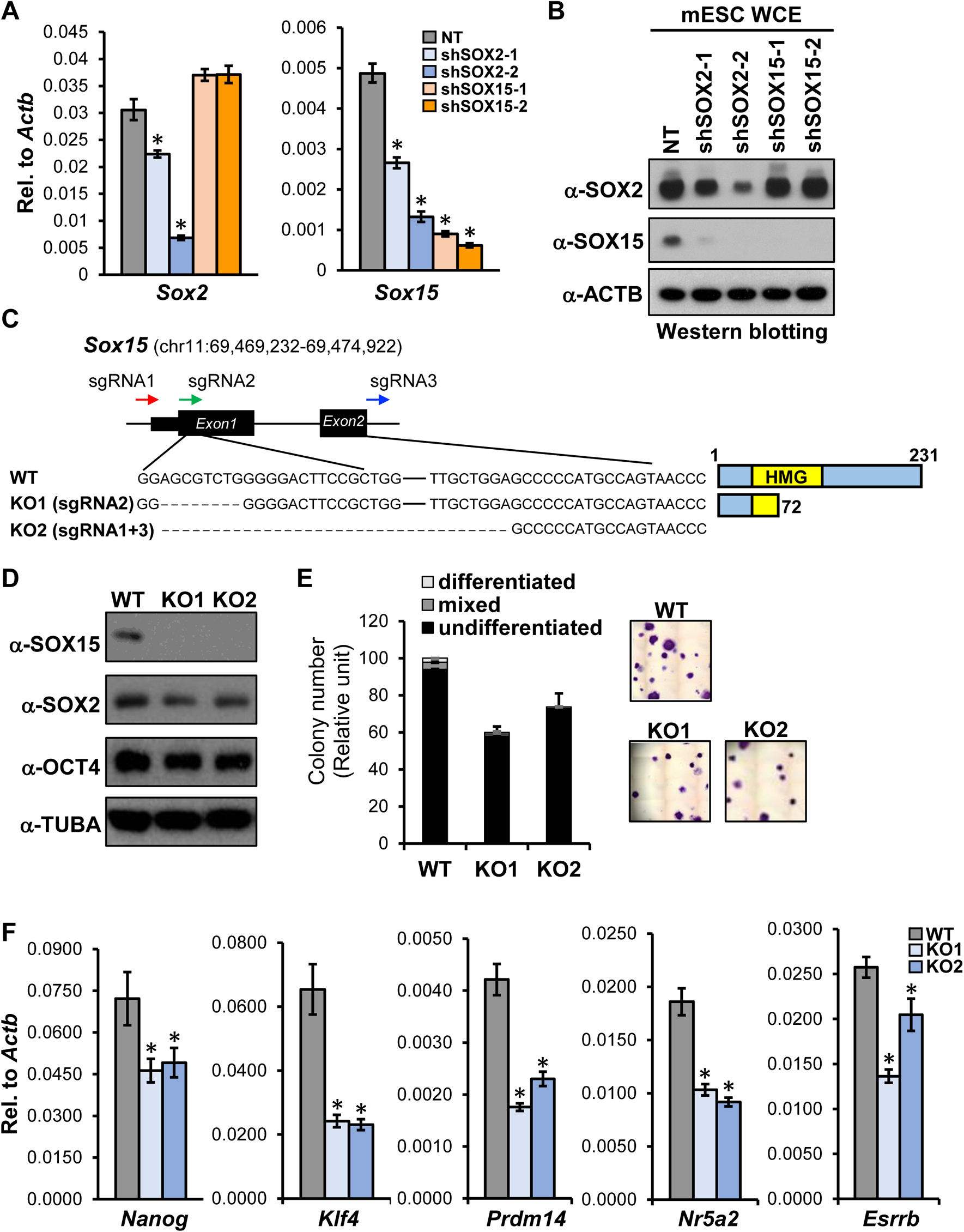
SOX15 deficiency compromises stem cell self-renewal and pluripotency gene expression. **(A)** Partial depletion of SOX2 by targeted shRNAs compromises SOX15 expression. Quantification of *Sox2* and *Sox15* mRNA levels in SOX2 or SOX15-knockdown ESCs are analyzed by qPCR and normalized to *Actb.* **(B)** SOX2 and SOX15 protein levels in WCEs of SOX2- and SOX15-knockdown ESCs are analyzed by western blotting, ACTB as loading control. **(C)** Left: Schematic diagram of *Sox15* deletion strategies using CRISPR/Cas9 system. K01 contains an 8 bp deletion induced by sgRNA2, leading to frameshift translation stop at amino acid (aa) 72 (full-length SOX15 contains 231 aa’s). K02 is generated by sgRNA1 and 3-mediated excision of the entire exon 1 and 2 of the *Sox15* gene. Right: Schematic representation of full­ length SOX15 in WT ESCs and predicted SOX15 fragment (1-72 aa) in K01. Relative position of HMG box is highlighted in yellow. **(D)** Western blotting of ESC WCEs confirms the absence of SOX15 in K01 and K02, while SOX2, OCT4, and a-tubulin (TUBA) levels remain largely the same as wild-type (WT) ESCs. **(E)** Colony formation assays of WT and *Sox15* KO ESCs. Differentiation status is evaluated on the basis of AP staining intensity and colony morphology. Representative images of AP staining of control and KO ESCs are shown (right). **(F)** Loss of SOX15 in ESCs compromises pluripotency gene expression. Quantification of mRNA levels of indicated pluripotency-associated genes by qPCR, normalized to *Actb.* Error bars represent SEM. *n* = 3. (*) *P* < 0.05, calculated by two-sided Student’s t-test.

ESCs are typically cultured in medium containing serum and the self-renewal cytokine leukemia inhibitory factor (LIF). Under this condition, ESCs exhibit a high degree of heterogeneity, reflected in their mosaic expression of several pluripotency-associated genes such as *Nanog* (36) and *Esrrb* (37). Cell populations expressing lower levels of these genes have reduced self-renewal capacity and are more prone to irreversible exit from pluripotency. However, this metastable pluripotent state can be stabilized in serum-free medium containing LIF and inhibitors (2i) of mitogen-activated protein kinase signaling and glycogen synthase kinase-3 (GSK3) (38, 39), which blocks signals that promote spontaneous differentiation. 2i/LIF conditions have been shown to bypass the requirement of TFs such as ESRRB and PRDM14 for self-renewal in serum/LIF (40, 41), suggesting that these TFs function to buttress the pluripotency gene network to counteract differentiation signals. Indeed, we found that 2i/LIF also overcame defects in self-renewal and restored expression of *Nanog* and *Prdm14* in *Sox15* KO ESCs (Figure S2C and S2D), indicating some SOX15-independent pathways enforced by 2i/LIF signaling may be sufficient to sustain the expression of these genes. Our results thus demonstrate that *Sox15* KO ESCs can be maintained in a stable pluripotent state under specific conditions, and that SOX15 stabilizes stem cell self-renewal in serum/LIF conditions by promoting optimal expression of key pluripotency genes such as *Nanog*, *Esrrb, and Prdm14*.

### Non-redundant function SOX15 in cellular reprogramming

Barriers to reacquisition of pluripotency during somatic cell reprogramming are high, which include the faithful reactivation of the pluripotency gene network in somatic cells (42–44). We reasoned that such processes may be more sensitive to perturbation of pluripotency-associated TFs. We therefore asked whether induced pluripotent stem cell (iPSC) generation requires SOX15 (18, 19). To address this question, we transduced mouse embryonic fibroblasts (MEFs) with lentiviruses expressing non-targeting control shRNA or two independent shRNAs specific for SOX15. We next initiated reprogramming of these MEFs by doxycycline-induced expression of OCT4, KLF4, SOX2, and c-MYC (45). Knockdown of SOX15 led to a marked decrease in the number of AP-positive iPSC colonies (Figure 3A). Flow cytometry analysis showed that loss of SOX15 interfered with the downregulation of fibroblast-associated cell surface marker THY1, and compromised the expression of iPSC early marker SSEA1 and the late stage marker EpCAM (Figure 3B) (46), which has been shown to correlate with the reactivation of *Nanog* and endogenous *Oct4* (47). Our results show that SOX15 is necessary for efficient suppression of somatic cell identity and reactivation of pluripotency-associated genes during reprogramming, and cannot be replaced by exogenously overexpressed SOX2.

**Figure 3.**
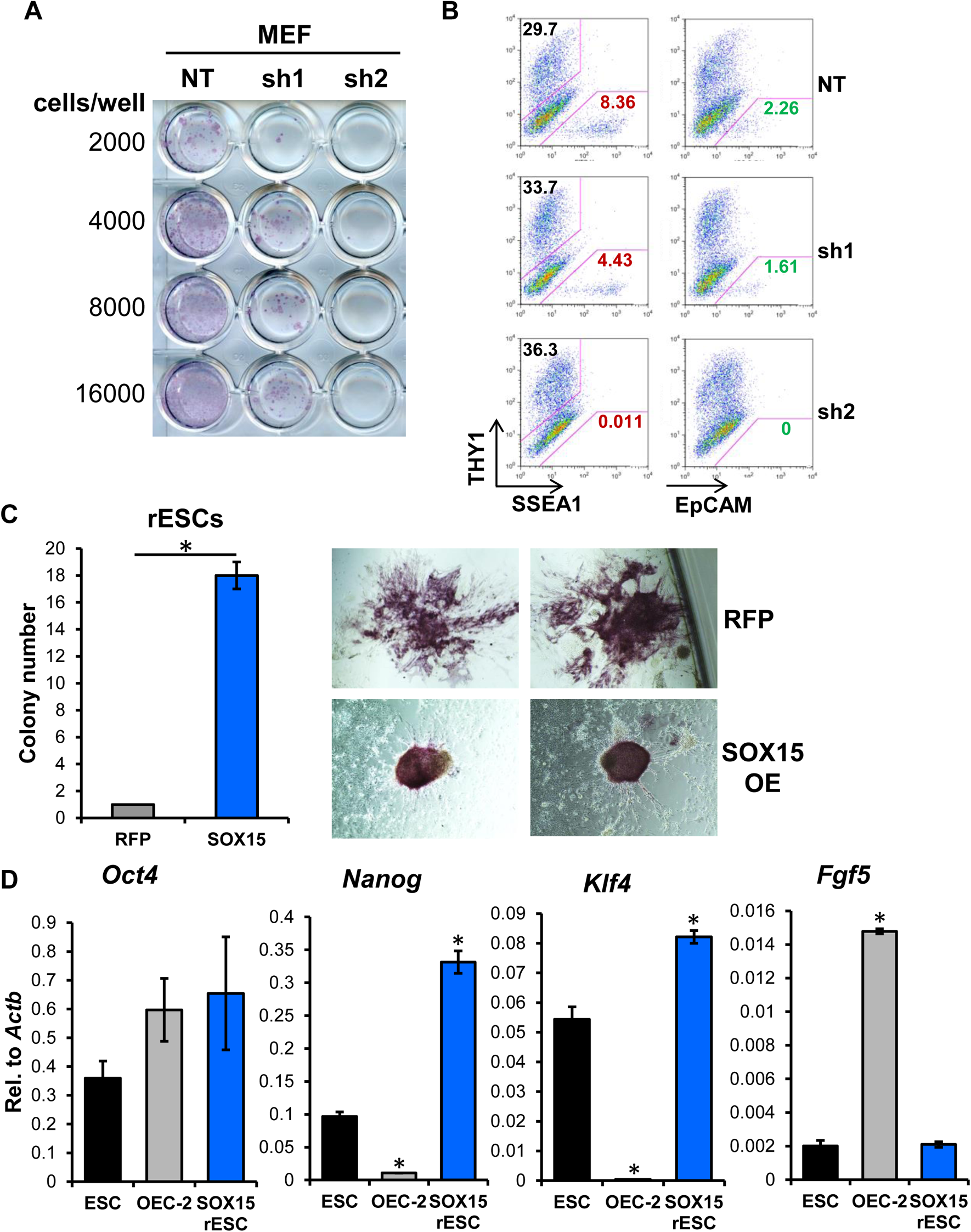
Regulation of cellular reprogramming by SOX15. **(A)** Depletion of SOX15 blocks iPSC generation from mouse embryonic fibroblasts (MEFs). Control (NT) and SOX15-knockdown (sh1 and 2) MEFs transduced with lentiviruses expressing reprogramming TFs (SOX2, OCT4, KLF4, and c-MYC) are plated at indicated cell numbers. Cells are stained for AP activity after 14 days post induction (dpi) of reprogramming TFs by doxycycline (dox) (11 days with dox followed by 3 days without dox). **(B)** Single cell suspensions of 14 dpi MEFs as described in (A) are stained with anti-THY1 together with anti-mouse SSEA1 or anti-EpCAM antibodies and analyzed by flow cytometry. Percentage of THY1+ and SSEA1+ cells are indicated. (C) OEC-2 epiblast stem cells (EpiSCs) are transduced with lentiviruses expressing SOX15 or RFP as control. Transduced cells are then replaced with 2i!LIF ESC medium. Colonies are stained for AP activity and counted after 14 days. Representative images of AP-staining of reprogrammed ESC (rESCs) colonies are shown (right). **(D)** mRNA levels of pluripotency-associated genes *(Oct4, Nanog,* and *Klf4)* and EpiSC-enriched marker *Fgf5* in ESCs, OEC-2, and SOX15-overexpressing rESCs are quantified by qPCR, normalized to *Actb.* Error bars represent SEM. *n* = 3. (*) *P* < 0.05, calculated by two­ sided Student’s t-test.

It is possible that the observed reprogramming block in SOX15 knockdown MEFs is due to defects in events prior to pluripotency gene network reactivation. To test more directly the role of SOX15 in induced pluripotency, we chose a different reprogramming paradigm, in which we assessed the ability of SOX15 to regulate the conversion between two related pluripotent states found in mouse ESCs and epiblast stem cells (EpiSCs). Unlike ESCs which are derived from pre-implantation blastocysts, EpiSCs are isolated from post-implantation embryos (48, 49). Similar to ESCs, EpiSCs express core TFs OCT4 and SOX2 and can self-renew and differentiate into three embryonic germ layers. However, in addition to changes in the OCT4/SOX2 transcriptional network in EpiSCs (50), EpiSCs require fibroblast growth factor (FGF) and Activin/Nodal signaling instead of LIF for self-renewal, and display restricted cell fates in part due to downregulation of several key pluripotency genes such as *Klf4* and *Nanog* and reactivation of lineage-specific genes representing the three germ layers (49, 51). EpiSCs are poised to undergo differentiation and thus represent a more advanced, “primed” pluripotent state as opposed to the more pristine, “naive” state found in ESCs. We found that SOX15 expression is restricted to ESCs when compared to an established EpiSC line, OEC-2 (52) while SOX2 protein level remains largely the same in both cell lines (Figure S3A and S3B). To ensure that the observed differences in SOX15 expression pattern is not a cell line specific phenomenon, we directly converted mouse ESCs to an EpiSC-like (EpiSC-L) state by culturing ESCs in medium containing FGF2 and Activin (52, 53) and confirmed that SOX15 expression was also markedly reduced in EpiSC-L cells (Figure S3C), concomitant with upregulation of lineage-specific genes such as *Fgf5* and *T* (Figure S3D). These observations prompted us to ask whether SOX15 could play a role in promoting the acquisition of naive pluripotency in EpiSCs.

By directly culturing EpiSCs in 2i/LIF medium, it has been shown that only a small fraction of EpiSCs are converted to ESC-like cells (rESCs) (52). However, this inefficient process can be enhanced by ectopic expression of naive pluripotency-enriched transcription factors such as KLF4 (52), ESRRB (40, 54), PRDM14 (55) and NANOG (56). Because we showed that expression of these genes is partially dependent on SOX15 in ESCs (Figure 2F), we next tested whether ectopic expression of SOX15 could also promote rESC generation. We transduced OEC-2 cells with lentiviruses expressing SOX15 or RFP as control (Figure S3E). We found that ectopic expression of SOX15 significantly increased reprogramming efficiency of EpiSC to rESC (Figure 3C). We further noticed that rESC colonies obtained from SOX15-overexpressing cells are tightly packed with more uniform AP staining, characteristics of *bona fide* ESCs (Figure 3C). By contrast, RFP-rESC colonies tended to be more loosely packed with large, elongated cells and variegated AP staining intensity, indicating that reprogramming may be less faithful than SOX15-rESCs. qPCR analysis confirmed SOX15-overexpressing rESCs restored the expression of key naive pluripotency-associated genes (*Nanog* and *Klf4*) and suppressed lineage-specific gene *Fgf5*, to levels comparable to ESCs (Figure 3D). Note that SOX2 level remains high in OEC-2 cells that is similar to ESCs (Figure S3B). Our results point to a role of SOX15 distinct from SOX2 in promoting the reacquisition of naive pluripotency in EpiSCs.

### *Sox15* deletion compromises stem cell pluripotency

To gain a more comprehensive view of the transcriptional programs regulated by SOX15, we performed RNA-seq analysis on three biological replicates each of wild-type (WT) and *Sox15* KO ESCs. ~7% of protein-coding genes in mouse ESCs are either up (830 genes) or down (589) regulated (Figure 4A). We performed unbiased gene ontology (GO) analysis on both up- and down-regulated transcripts and observed a significant overrepresentation of categories related to stem cell maintenance and blastocyst formation, consistent with a role of SOX15 in pluripotency (Figure 4B). Interestingly, GO analysis also identified genes involved in cell migration, trophectodermal specification, and neural stem cell and neuronal differentiation. Gene transcripts associated with immune responses, ion transport, and cell-cell adhesion are elevated in *Sox15* KO ESCs (Figure S4). In agreement with our qPCR results (Figure 2F), key pluripotency genes such as *Nanog*, *Klf4*, and *Esrrb* are downregulated in *Sox15* KO ESCs (Figure 4C). However, genes related to trophectoderm and neuronal differentiation were also decreased compared to WT ESCs. This result was somewhat unexpected given that *Sox15* KO ESCs are more prone to spontaneous exit from pluripotency, and thus one might anticipate that lineage-specific genes should be upregulated compared to WT ESCs. However, SOX15 has previously been implicated in trophectoderm differentiation by cooperating with TF HAND1 (57). While the role of SOX15 in neurogenesis has not been reported, we found that neurogenic transcription factors such as *Sall3* (58, 59) and *Nfib* (60), and Slit/Robo signaling, previously shown to modulate neurogenesis (61, 62), are downregulated in *Sox15* KO ESCs. Reduced levels of these transcripts in ESCs may reflect that, as *Sox15* KO ESCs transiently exit from pluripotency under serum/LIF condition, these cells may have a reduced capacity to commit towards the neural cell fates by upregulating neurogenic gene expression programs. These observations suggest that *Sox15* KO ESCs may have restricted potential to differentiate into the three embryonic germ layers.

**Figure 4.**
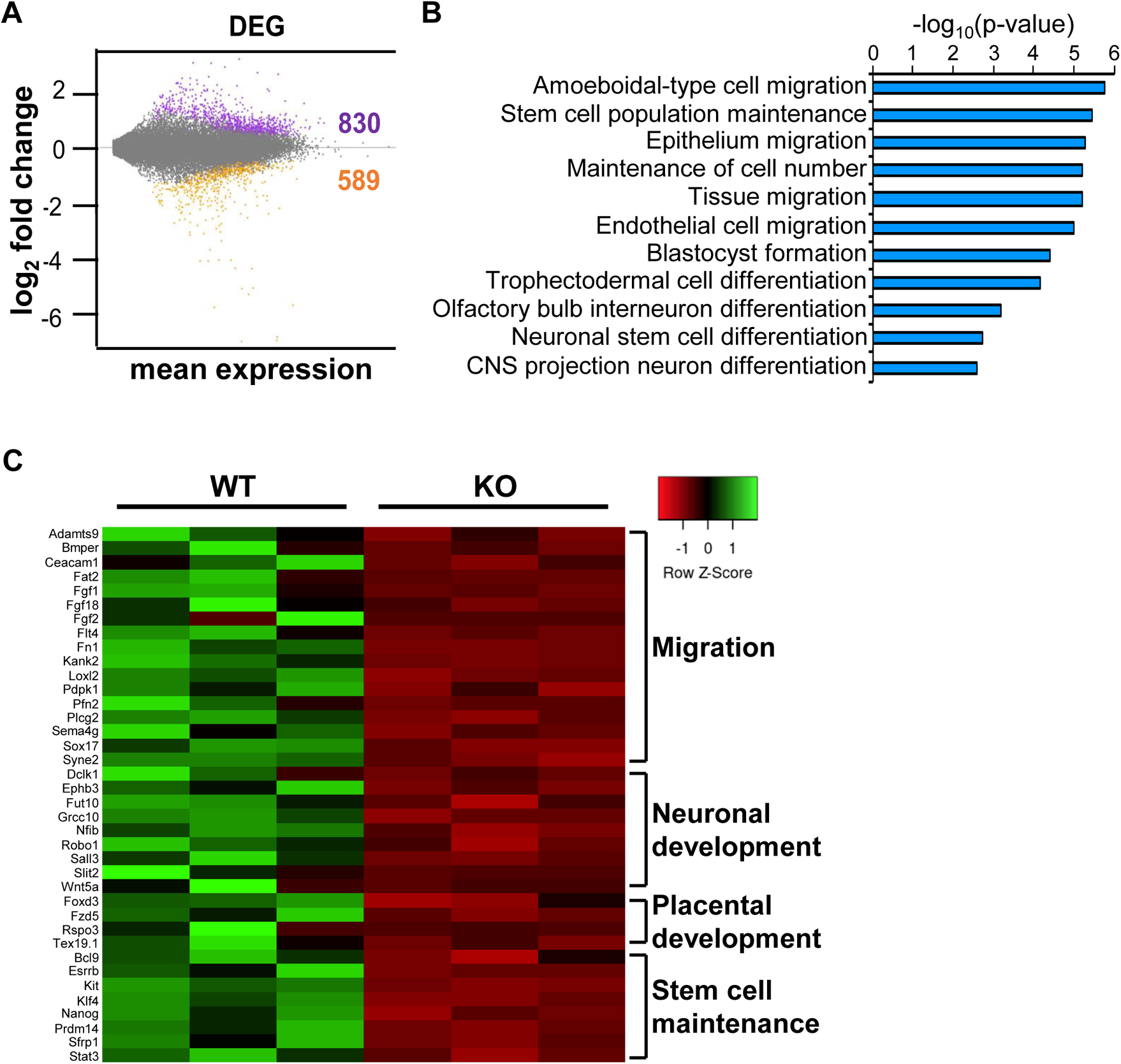
Gene expression profile of Sox15-knockout ESCs. **(A)** RNA-seq analysis identifies differentially expressed genes (DEGs) in *Sox15* K01 ESCs compared to VVT, plotted as a function of mean expression (fpkm) and fold changes (log2). Number of genes upregulated (830) and downregulated (589) are indicated. **(B)** Gene ontology (GO) analysis on DEGs in *Sox15* K01 ESCs compared with all annotated genes. p-value per GO classification is shown (-log1*O(p­* value)). **(C)** Heatmap showing the relative expression levels of genes indicated from three biological replicates each of VVT and *Sox15* KO ESCs. Scaled values indicate relative downregulation (red) or upregulation (green) of gene expression. Genes from each category are manually curated.

### SOX15 is required for *Hes5* expression and neural differentiation

Using ChIP-seq with an anti-HA antibody against HA-tagged SOX15 in our knock-in mouse ESC line, we identified a SOX15-bound region ~17 kb upstream of the *Hes5* gene locus (Figure 5A). *Hes5* belongs to the *Hairy and Enhancer of Split* (*Hes*) family transcriptional repressors that include *Hes1* and *Hes3* (63). *Hes1* and *Hes5* are expressed in the ventricular zone of the developing brain, in which neural stem cells (NSCs) reside (64, 65). In mice, it has been shown that *Hes1* and *Hes5* display some degree of functional redundancy in promoting self-renewal and maintenance of NSCs (65, 66). However, these repressors appear to also regulate the timing of cell fate specification of neural progenitor cells derived from NSCs (67–69). Therefore, the *Hes* family transcriptional repressors play complex roles in controlling both the maintenance of NSCs and their differentiation during development. SOX2 has been implicated in regulating *Hes5* expression in neural progenitor cells by binding at promoter proximal regions (70, 71). However, the role of SOX15 in neural differentiation is unknown.

**Figure 5.**
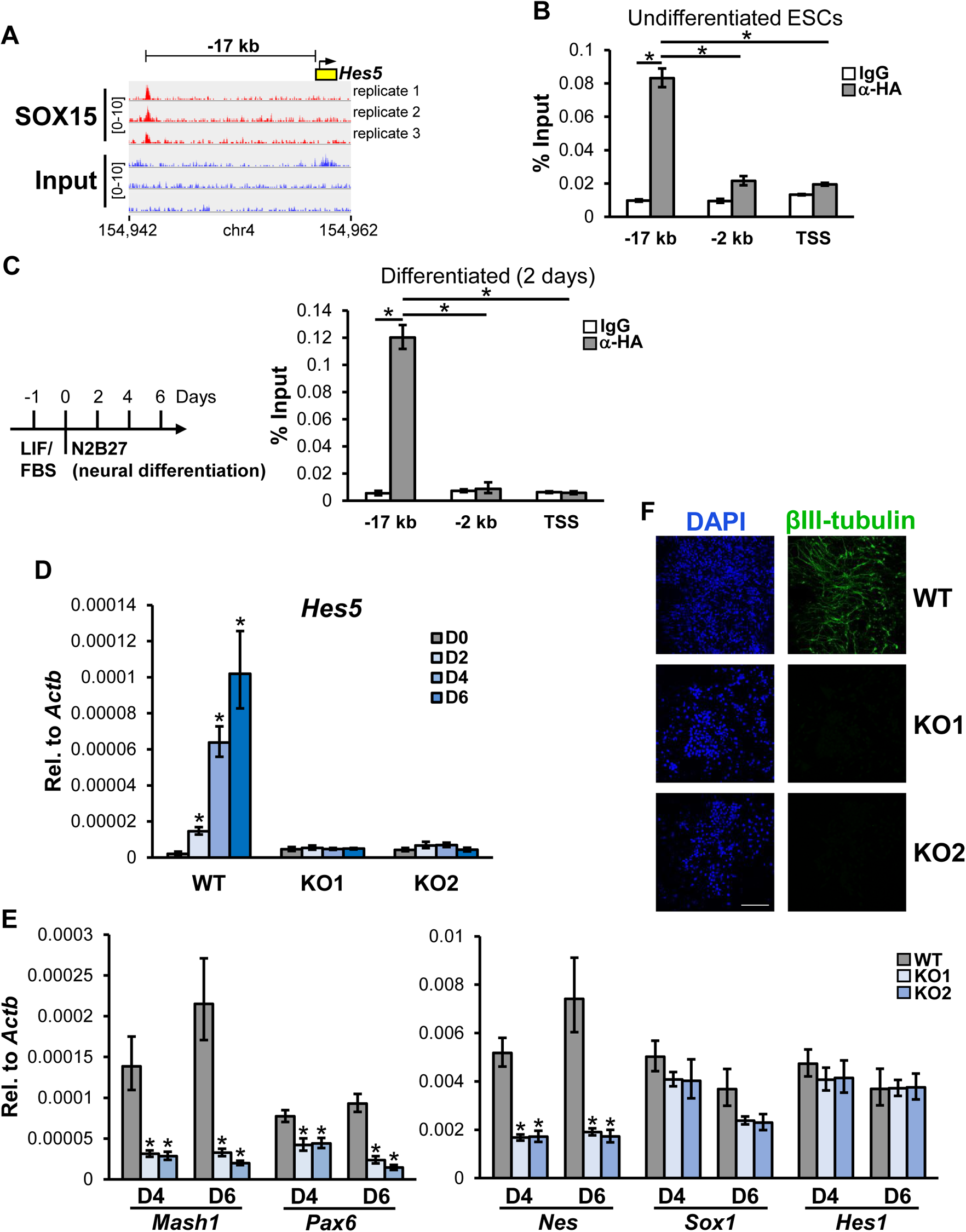
SOX15 regulates Hes5 expression and neural differentiation. **(A)** Integrative Genomics Viewer (IGV)-computed ChiP-seq tracks from three biological replicates are plotted as (number of reads) x [1,000,000/(total read count)] for the *Hes5* gene locus. Reads from SOX15-HA ChiP DNA and input chromatin are shown. Genomic coordinates are in kilobases. **(B)** MNase­ ChiP analysis of SOX15 occupancy at −17 kb, −2 kb, and TSS of the *Hes5* gene in undifferentiated ESCs. Enrichment and data analysis are as described in Figure 1C. **(C)** Schematic diagram describing differentiation of ESCs into neuroectodermal precursors. ESCs cultured in LlFifetal bovine serum (FBS) are plated 1 day before replacing with serum-free N2B27 differentiation medium. Cells are collected at 2, 4, and 6 days for analysis (left). MNase-ChiP analysis of SOX15 occupancy at the *Hes5* locus in differentiating ESCs (2 days), analyzed as described in (B). **(D)** *Hes5* mRNA levels in undifferentiated (DO) and differentiating VVT, *Sox15* K01 and K02 ESCs collected at day 2, 4, and 6 of differentiation, analyzed by qPCR and normalized to *Actb.* **(E)** mRNA levels of indicated neural markers in differentiating VVT and KO ESCs (D4 and 6) are quantified by qPCR and normalized to *Actb.* **(F)** Representative immunofluorescence images of VVT and *Sox15* KO (K01 and K02) ESCs after 6 days of differentiation, stained with antibodies against neuronal marker J3lll-tubulin. Nuclei are stained with DAPI. Scale bar, 100 IJ-m. Error bars represent SEM. *n* = 3. (*) *P* < 0.05, calculated by two-sided Student’s t-test.

Although the ChIP-seq assay yielded a limited number of binding sites, we confirmed binding using MNase ChIP-qPCR and demonstrated specific enrichment of SOX15 at −17 kb but not −2 kb or TSS of the *Hes5* locus in undifferentiated ESCs (Figure 5B and figure S5A and S5B). To examine the potential role of SOX15 in regulating *Hes5* expression during neural differentiation, we directly differentiated ESCs in serum-free, pro-neurogenic medium (72). *Hes5* expression was induced in SOX15-HA knockin ESCs 2 days after differentiation (Figure S5C). Even though SOX15 mRNA and particularly protein levels were downregulated at day 2 (Figure S5D), we found that SOX15 remained engaged at the −17 kb region, with no change in relative enrichment at −2 kb or TSS (Figure 5C). These results suggest that SOX15 is already bound at the −17 kb region prior to *Hes*5 induction during neural differentiation.

To examine whether SOX15 is required for *Hes5* expression, we compared its expression in WT and *Sox15* KO ESCs during neural differentiation. We observed robust induction of *Hes5* in WT ESCs over the course of 6-day differentiation process, wherein a population of neuroectodermal precursors such as NSCs and neural progenitors are formed (Figure 5D) (72). However, both *Sox15* KO ESC lines completely failed to induce *Hes5* expression. Consistent with the function of HES5 in NSC maintenance and multipotency (68), we also found that expression of NSC marker such as *Nestin* (*Nes*) and transcription factors important for neural commitment, *Mash1* and *Pax6*, were significantly compromised in *Sox15* KO ESCs, while *Sox1* was moderately reduced at day 6 (Figure 5E). Interestingly, even though *Hes1* expression was not affected in *Sox15* KO ESCs, we observed a pronounced defect of KO ESCs in expressing neural marker βIII-tubulin (Tuj1) (Figure 5F) (72), indicating that loss of *Hes5* expression cannot be compensated by *Hes1*. Given that SOX2 levels remained the same in WT and *Sox15* KO ESCs and during the early stages of differentiation (Figure S5D), our results uncovered a critical function of SOX15 in regulating *Hes5* expression and neural differentiation in vitro.

The −17 kb region bound by SOX15 displays distal enhancer signatures such as DNase I hypersensitivity and enrichment of H3K4 trimethylation (H3K4me3) and H3K27 mono-acetylation (H3K27Ac) (ENCODE) (Figure S6A). This region is also surrounded by several genes including *Tnfrsf14*, *Pank4*, and *Plch2* (Figure S6B). While there are no reports that directly implicate these genes in early neural fate specification, we tested whether expression of these genes is also dependent on SOX15. We found that expression of *Tnfrsf14*, *Pank4*, and *Plch2* in undifferentiated (D0) and differentiated ESCs (D6) was not affected by SOX15 depletion (Figure S6C). Therefore, it appears that the −17 kb region occupied by SOX15 may be specifically required for *Hes5* expression. It has previously been shown that RNA polymerase II (Pol II) recruitment to enhancers correlate with enhancer activities and gene activation (73). Therefore, to evaluate the activity of this putative distal enhancer in WT and *Sox15* KO ESCs at the onset of neural differentiation, we examined Pol II occupancy at the −17 kb region at day 2 of differentiation. We found striking reduction of Pol II occupancy at the −17 kb region in KO ESCs compared to WT, as well as at the −2 kb region and TSS (Figure 6A). These results are consistent with a defect in activating the −17 kb enhancer and transcription of *Hes5* in *Sox15* KO ESCs during neural differentiation. Importantly, SOX2 binding at the *Hes5* locus is not significantly affected by the absence of SOX15 in KO ESCs (Figure S6D). Together, our data point to a specific requirement of SOX15 in regulating enhancer and promoter activities of *Hes5* during early stages of neural differentiation.

**Figure 6.**
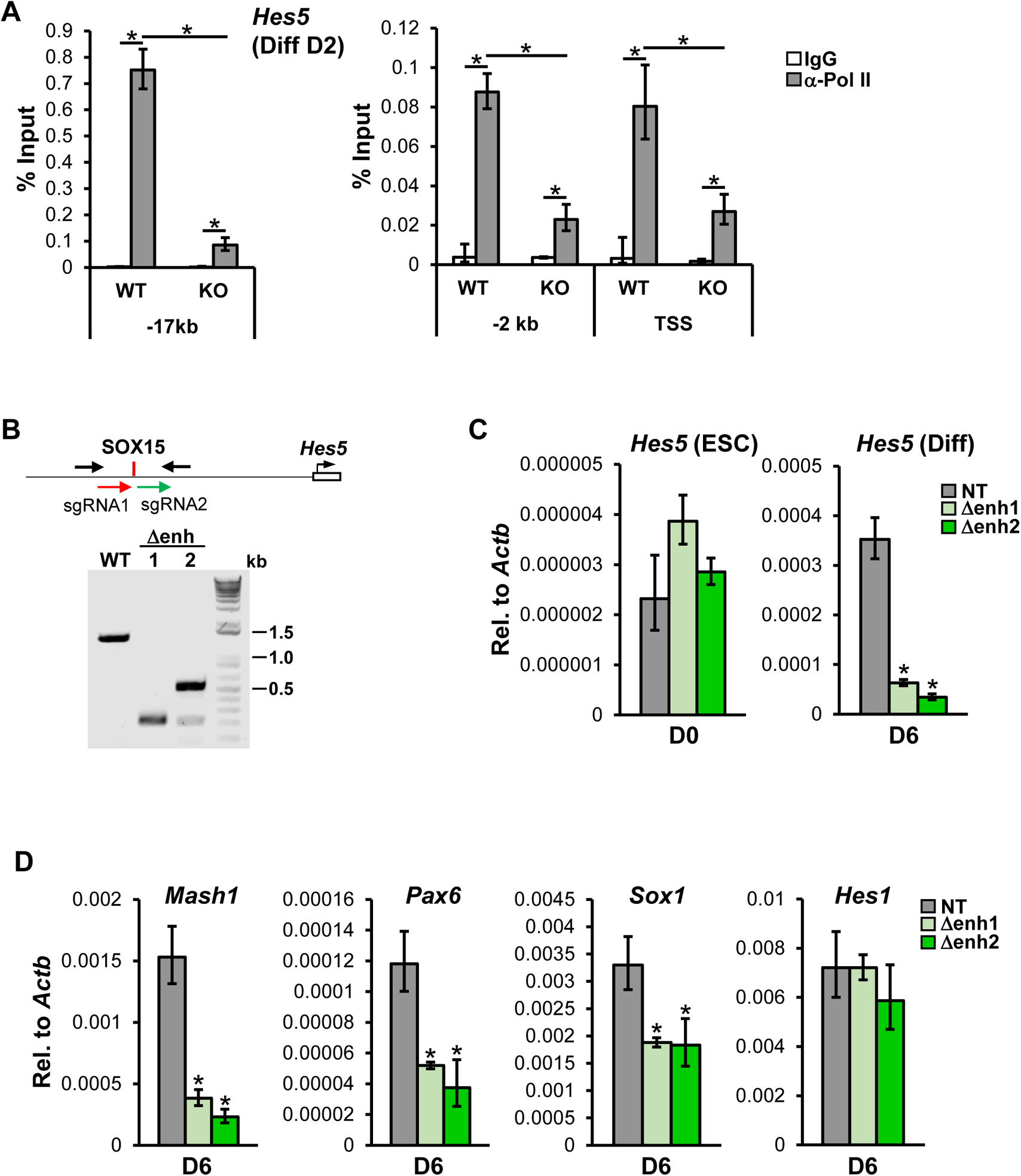
SOX15-bound −17 kb region controls Hes5 gene expression during neural differentiation. **(A)** MNase-ChiP analysis of Pol II occupancy at indicated regions of the *Hes5* gene locus in WT and *Sox15* knockout (K02) ESCs after 2 days of differentiation (Diff 02). Representative data showing the enrichment of Pol II (gray bars) compared to control lgGs (white bars) are analyzed as described in Figure 1C. **(B)** Schematic diagram showing CRISPR/Cas9 mediated deletion of the −17 kb region using two sgRNAs flanking a −1 kb region (top). PCR analysis of genomic DNA from WT and two independent homozygous deletion ESC clones (t.enh1 and 2) (bottom). Primers used to amplify the deleted region indicated (black arrows). **(C)** *Hes5* mRNA levels in undifferentiated WT and t.enh ESCs and their differentiated (day 6, 06) counterparts are analyzed by qPCR, normalized to *Actb.* **(D)** mRNA levels of neural markers *(Mash1, Pax6, Sox1,* and *Hes1)* in differentiated WT and t.enh ESCs (06) are quantified by qPCR, normalized to *Actb.* Error bars represent SEM. *n* = 3. (*) *P* < 0.05, calculated by two-sided Student’s t-test.

To more firmly establish the −17 kb region as an important enhancer that drives *Hes5* expression during neural differentiation, we employed CRISPR/Cas9 gene editing to delete this putative enhancer. Two independent homozygous deletion ESC lines (Δenh1 and 2) were generated and confirmed by genomic DNA PCR and sequencing (Figure 6B). Deletion of this enhancer region did not affect the expression of core pluripotency genes (*Nanog*, *Oct4*, and *Sox2*), genes in the vicinity of the −17 kb region (i.e., *Tnfrsf14*, *Pank4*, and *Plch2*), or basal expression of *Hes5* in undifferentiated ESCs (Figure 6C and figure S6E and S6F). However, compared to control ESCs expressing a non-targeting sgRNA, induction of *Hes5* in Δenh1/2 was significantly compromised at day 6 of differentiation while *Hes1* mRNA levels were unaffected (Figures 6C and 6D). *Mash1*, *Pax6*, and *Sox1* were also reduced in Δenh1/2 compared to control ESCs, indicating that neural differentiation was compromised (Figure 6D). These results are reminiscent of the differentiation defect observed in *Sox15* KO ESCs (Figures 5E and 5F), suggesting that activation of *Hes5* by SOX15 is an important early event in neural differentiation in vitro. It has recently been reported that *Hes5* is a direct target of the TF GLI3 in the Hedgehog (Hh) pathway, which plays an important role in neural differentiation during development (74). We found that *Gli3* mRNAs were elevated in WT ESCs during neural differentiation (Figure S6G). However, no significant differences in *Gli3* levels were observed between WT and KOs at day 4 of differentiation (Figure S6H), even though *Hes5* expression was already reduced in *Sox15* KO cells (Figure 5D). Therefore, *Gli3* alone does not appear to be sufficient to induce *Hes5* expression. Using in vitro neural differentiation system as a paradigm for studying the role of Sox TFs in regulating exit from pluripotency and lineage specification, we uncovered an important function of SOX15 in the early neural fate commitment that appears to be distinct from SOX2.

## Discussion

### SOX15 function in maintenance and reacquisition of pluripotency

In this study, we show that SOX15 and SOX2 activate transcription of pluripotency genes due to their ability to recruit a common set of transcriptional coactivators enriched in ESCs. Outside the highly conserved HMG domain (~80% sequence identity), SOX15 and SOX2 protein sequences diverge significantly, raising questions on how SOX15 and SOX2 (but not other Sox family TFs) are able to stimulate transcription using the same transcription factor ensemble. We have recently shown that one of the stem cell-enriched coactivators, ABCF1, activates ESC-specific transcription by selectively interacting with the HMG domain of SOX2 (but not SOX11, for example) (15). We find here that ABCF1 also binds SOX15, thus revealing a role of cell-specific coactivators in discriminating various SOX-OCT4 complexes that may assemble at a gene promoter in ESCs, likely by recognizing small differences in the HMG box sequence. Taken together with our previous observation that OCT4 alone with SCCs is not sufficient to activate *NANOG* transcription (12), our results indicate that the ability of SOX15 to bind ABCF1 is a critical determinant for transactivation. However, considering that SOX15 abundance in ESCs is significantly lower than that of SOX2, disruption of a SOX15-ABCF1 interaction alone may not be sufficient to account for the degree with which genes targeted by both SOX15 and SOX2 are downregulated in *Sox15* KO ESCs. These observations suggest non-redundant mechanisms by which SOX15 contributes to pluripotency gene activation. Sox TFs have been shown to interact with various chromatin remodeling and modifying complexes to drive cell type-specific transcription (71, 75–77). It is possible that the unique C-terminal region outside of the HMG box also play a role, potentially by mediating SOX15-specific interactions that are conducive to gene activation in the context of chromatin in ESCs. Furthermore, studies have shown that the unique region can modulate DNA binding specificity of the HMG domain (78). Future studies will be required to examine how these SOX15-dependent interactions may contribute to stem cell self-renewal and pluripotency gene expression.

*Sox15*-null mice are viable, suggesting no overt defects in the formation of pluripotent cells in the inner cell mass (ICM) of the blastocyst (22). Given the shared mechanisms by which SOX15 and SOX2 regulate the expression of core pluripotency genes, SOX2 alone may be sufficient to initiate pluripotency in the ICM, likely because the pluripotent state in ICM is relatively transient during development. By contrast, our data suggest that long-term maintenance of ESCs in serum/LIF requires the combined action of SOX2 and SOX15, likely by counteracting cell-intrinsic and extrinsic signals that promote ESCs to exit from pluripotency. Our study also reveals a critical role of SOX15 in overcoming barriers to reacquisition of pluripotency during reprogramming. It is known that several key primordial germ cell (PGC) TFs including PRDM14 promote iPSC generation (55, 79–81). Coupled with recent reports implicating a role of SOX15 in PGC fate specification (82, 83) and our observation that *Prdm14* is downregulated in *Sox15* KO ESCs, we speculate that SOX15 may regulate reprogramming by promoting the expression of critical PGC-associated TFs. It is also possible that SOX15, together with SOX2, ensures optimal reactivation of the pluripotency gene network during reprogramming. Paradoxically, ectopic expression of SOX15 in place of SOX2 dramatically reduces reprogramming efficiency in one study (21) or fails to generate iPSCs from fibroblasts altogether (84). However, exogenous SOX15 can promote iPSC derivation as efficiently as SOX2 from a cell type that is considered more permissive to TF-induced reprogramming (e.g., menstrual blood stromal cells (MnSCs)) (84) These observations indicate that SOX15 has an intrinsic capacity to induce pluripotency, consistent with our data showing that it can activate the transcription of key pluripotency genes by partnering with OCT4 and SCCs. However, it appears that this activity is cell-context dependent. We surmise that the epigenetic environment and transcription factor repertoire in MnSCs that allow SOX15 to efficiently target and activate pluripotency genes may be less favorable or underdeveloped in skin-derived fibroblasts. This may also explain why endogenous SOX15 is still required for iPSC generation from fibroblasts when SOX2 is already overexpressed. As fibroblasts traverse through various reprogramming intermediate stages, the accompanied changes in chromatin landscape and coactivator expression levels may become more permissive to activation of SOX15 target genes important for pluripotency reacquisition (85). Similarly, we posit that during EpiSC to ESC conversion, the more relaxed, “open” chromatin environment associated with pluripotent EpiSCs (compared to fibroblasts) may facilitate transcriptional activation of naive pluripotency-associated genes by exogenous SOX15 (86). Human ESCs have been shown to share features with mouse EpiSCs including their restricted differentiation potential, which in turn limit the use of human ESCs for regenerative medicine and human disease modeling (87, 88). The ability of SOX15 to promote an ESC-like state in mouse EpiSCs may inform new strategies to promote naive pluripotency in human ESCs.

### SOX15 function in neural differentiation

We noticed that SOX15 is already engaged at the −17 kb distal enhancer of the *Hes5* gene in undifferentiated ESCs. This observation suggests that SOX15 is necessary but not sufficient for *Hes5* induction. Because SOX2 recruitment to the *Hes5* locus does not appear to be affected by the absence of SOX15 in KO ESCs, we propose that transcriptional activation of *Hes5* by SOX15 likely requires other TFs and coactivators that are induced at the onset of neural differentiation, in conjunction with epigenetic remodeling at the *Hes5* gene locus (89). The pre-binding of SOX15 at the *Hes5* locus may represent a common strategy by which developmentally poised genes are regulated upon differentiation (90).

*Sox15* knockout mice appear phenotypically normal, indicating no gross defect in neurogenesis (23, 91). Interestingly, we find that *Hes1* expression in ESCs is not affected by the absence of SOX15. Given that multiple *Hes* genes appear to be functionally redundant in development (64, 92, 93), it is conceivable that normal *Hes1* expression levels in *Sox15* KO mice may be sufficient to support neurogenesis. However, whether some neurological changes exist in *Sox15* KO mice remains to be investigated. While the precise mechanism of mammalian neurogenesis remains unclear (94), at least in vitro, our results suggest a hierarchical organization of HES proteins, in which *Hes5* appears to be play an important role in promoting the commitment of ESCs to NSC fate and neural lineage. This is in agreement with previous observations wherein *Hes5* expression correlates with NSC identity and multipotency better than *Hes1* (68). However, other neurogenic TFs and signaling pathways that are downregulated in *Sox15* KO ESCs (Figure 4C) may also contribute to this process. We show that SOX15 levels decrease sharply shortly after neural differentiation is initiated. We hypothesize that this may allow other Sox TFs including SOX2 to precisely tune *Hes5* expression levels in a spatiotemporal manner to control NSC maintenance vs neural cell fate specification (70, 95, 96). In summary, our findings reveal a multifaceted role of SOX15 in regulating the pluripotent stem cell fate and execution of neural differentiation programs.

## Experimental Procedures

### DNA constructs and antibodies

Complementary DNAs (cDNAs) for human OCT4, SOX2, and SOX15 were obtained from cDNA libraries generated from human embryonic carcinoma cell line NTERA-2 (NT2). Intronless human SOX1 and SOX11 cDNAs were amplified from human genomic DNA. C-terminal FLAG-tagged OCT4 and various SOX TFs were generated by subcloning cDNAs into pFLAG-CMV5a mammalian expression vector (Sigma-Aldrich). For stable expression of mCherry and SOX15 in OEC-2 cells, mCherry and SOX15 cDNA were cloned into lentiviral overexpression vector pHAGE-IRES-Neo (12). Untagged human DKC1 cDNA was subcloned into pHAGE-IRES-Neo. Mammalian expression constructs for untagged XPC-RAD23B-CETN2 complex (pHAGE-XPC.com) (12) and V5-tagged ABCF1 (pFLAG-CMV5a-ABCF1-V5) (15) were described. Note that the V5 tag followed by a translation stop codon was inserted upstream of the built-in C-terminal FLAG tag in pFLAG-CMV5a vector. pLKO lentiviral vectors expressing non-targeting shRNA and shRNAs targeting SOX2 and SOX15 were purchased from Sigma-Aldrich.

Polyclonal antibodies against SOX2 were purchased from EMD Millipore (AB5603); XPC (A301-122A) from Bethyl Laboratories; DKC1 (H-300) and OCT4 (N-19) from Santa Cruz Biotechnology; SOX15 (25415-1-AP) from ProteinTech. Monoclonal antibodies against hemagglutinin (HA) tag (C29F4) for ChIP and western blotting were purchased from Cell Signaling Technology; ACTB (66009-1) from ProteinTech; FLAG tag (M2) and α-tubulin (T5168) from Sigma-Aldrich; βIII-tubulin from Biolegend (801201).

### Cell culture

NT2 cell line was obtained from the American Type Culture Collection (ATCC). NT2 and 293T cells were cultured in Dulbecco’s modified Eagle’s medium (DMEM) high glucose with GlutaMAX (2 mM; Gibco) supplemented with 10% fetal bovine serum (FBS, Gibco). Large scale culture of NT2 cells to generate P1M fraction was described (12).

D3 mouse ESC line was purchased from ATCC and adapted to feeder-free condition as described (12). Medium was changed daily. Cell cultures were passaged by trypsin. For ChIP-seq experiments, SOX2-HA and SOX15-HA knockin mouse ESC lines were first adapted to 2i/LIF culture condition as described (15). Differentiation of ESCs by embryoid body (EB) formation was performed as described (97). EBs were formed by using the hanging drop method. Briefly, hanging drops (30 μl) containing ~300 cells were pooled after 2 days and transferred to low attachment plates. EBs were collected for analysis after 8 days of differentiation. Differentiation of ESCs into neuroectodermal precursors was performed as described (72). Differentiation medium was changed every other day for 6 days. EpiSC line OEC-2 was kindly provided by Dr. Austin Smith laboratory. OEC-2 was cultured on human fibronectin-coated plates (15 μg/ml; EMD Millipore) in serum-free N2B27 medium (DMEM/F12 [Gibco] and Neurobasal [Gibco] 1:1 ratio, 1:100 N2 Supplement [Gibco], 1:50 B27 Supplement [Gibco], 2 mM Glutamax [Gibco], beta-mercaptoethanol [0.1 mM, Sigma-Aldrich], and 1x penicillin and streptomycin [100 U/ml, Gibco]) supplemented with 20% KnockOut Serum Replacement (Gibco), bovine serum albumin (0.01%, Sigma-Aldrich), Activin A (20 ng/ml, R&D Systems), and FGF2 (12 ng/ml, R&D Systems). For in vitro differentiation of EpiSC-L cells. ESCs cultured on fibronectin-coated plates in 2i/LIF medium were exchanged with serum-free N2B27 with Activin A and FGF2 as described above with the addition of 1% KnockOut Serum Replacement (Gibco). EpiSC-L cells are collected for analysis after 10 days of differentiation.

### In vitro transcription assay

The human *NANOG* transcription template, purification of activators OCT4 and SOX TFs, general transcription factors, RNA polymerase II, recombinant XPC complex and DKC1 complexes reconstituted from insect Sf9 cells (12, 13), and recombinant ABCF1 purified from *Escherichia coli* (15) were described. Partially-purified phosphocelluose 1M (P1M) nuclear cell extracts from NT2 cells were described (12). In vitro transcription and primer extension assays were described (12).

### Coimmunoprecipitation assay

pHAGE-XPC.com plasmid expressing XPC and its subunits RAD23B and CETN2, pHAGE-DKC1-IRES-Neo, pFLAG-CMV5a-ABCF1-V5 were co-transfected with control empty pFLAG-CMV5a vector or pFLAG-CMV5a-SOX15 into 293T using polyethylenimine (PEI, Polysciences) at a DNA:PEI (μg) ratio of 1:2.5. 40 hours after transfection, cells on 10-cm dishes were lysed with 1 ml of lysis buffer [50 mM HEPES pH 7.6, 0.12 M NaCl, 1% NP-40, 1 mM EDTA, and 10% glycerol] containing 1 mM DTT, 0.25 mM PMSF, and complete protease inhibitor cocktail (Sigma-Aldrich). Cell lysates were cleared by centrifugation at 15,000 rpm for 30 min at 4°C. Cleared lysates were then incubated with anti-FLAG M2 agarose for 3 to 4 hours at 4°C. Bound proteins were washed extensively with lysis buffer followed by FLAG peptide elution (0.4 mg/ml) in elution buffer [50 mM HEPES pH 7.6, 0.15 M NaCl, 0.5% NP-40, 10% glycerol) containing 0.5 mM DTT, 1 mM benzamidine, and 0.5 mM PMSF.

### Lentiviral production and transduction

For lentivirus production, nontargeting control and pLKO plasmids targeting mouse SOX15 (Sigma-Aldrich), or pHAGE plasmids for overexpression of SOX15 or control mCherry were cotransfected with packaging vectors (psPAX2 and pMD2.G; Addgene) into 293T cells using PEI methods. Supernatants were collected at 48 hours and again at 72 hours. Virus preparation was performed as described (12). Functional viral titer was determined by transduction of limited dilution of concentrated viruses into HeLa or NIH 3T3 cells followed by counting antibiotic-resistant colonies to obtain colony-forming unit per milliliter. Cells were infected with viruses in the presence of polybrene (8 μg/ml). For knockdown experiments, mouse ESCs transduced with pLKO viruses (multiplicity of infection [MOI] of 5) were selected with puromycin (1.5 μg/ml). Stable expression of SOX15 or mCherry in transduced OEC-2 cells was maintained in serum-free N2B27 medium containing geneticin (500 μg/ml).

### Cellular reprogramming and flow cytometry

For generation of iPSCs from fibroblasts, CF-1 MEFs (Charles River Laboratories) were transduced with inducible STEMCCA and rtTA lentivirus–containing supernatants overnight in polybrene (8 μg/ml; Sigma-Aldrich). Cells were expanded and transduced with pLKO viruses expressing control shRNA or shRNAs targeting SOX15. Doxycycline (2 μg/ml; Sigma-Aldrich) was supplemented to ESC medium to induce expression of OCT4, KLF4, SOX2, and c-MYC. Reprogramming was assayed by AP staining (EMD Millipore) or by flow cytometry analysis using anti-SSEA1, anti-THY1, and anti-EpCAM (BioLegend) on BD LSRFortessa, performed according to the manufacturers’ protocols.

For EpiSC to ESC conversion, ~1×10^6^ stable OEC-2 cells expressing SOX15 or mCherry (MOI of 3) were plated in OEC-2 culture medium. After 24 hours, medium was changed to 2i/LIF medium. 2i/LIF medium was changed every other day for 14 days.

### ChIP-seq analysis

HA-ChIP assays were performed in SOX15-HA knockin mouse ESCs essentially as described (13) except cells were cross-linked with formaldehyde (1%) for 5 min and cross-linked chromatin immunoprecipitated using an anti-HA rabbit monoclonal antibody (Cell Signaling Technology). Knock-in ESCs were first adapted to serum-free 2i/LIF condition as described before ChIP (15). ChIP-seq DNA libraries were constructed using the KAPA HyperPlus DNA Library Prep Kit for Illumina (Roche) according to the manufacturer instructions. Purified libraries were sequenced on NextSeq Illumina sequencer using NextSeq.

ChIP-seq data was processed using the ChIP-seq pipeline implemented in bcbio-nextgen v1.2.0.(https://bcbio-nextgen.readthedocs.org/). Briefly, raw sequence quality was evaluated using FASTQC (98). Reads were filtered and trimmed with Atropos (99) and mapped to the mouse genome build mm10 using Bowtie2 (100) with default parameters. Peaks were called using MACS2 (101) with default parameters after excluding multi-mapping reads and removing duplicates. Quality control was performed with ChIPQC (102) and DeepTools (103) to assess peak quality, coverage and reproducibility of the peaks across replicates. Likely peak artifacts were filtered out using the ENCODE blacklist (104). Overlapping peaks were identified using Bedtools (105). ChIP-seq peaks were annotated and functional enrichment analysis was performed using ChIPseeker (106). Known and de novo motifs were identified with HOMER (107). Data visualization was performed using the Integrated Genomics Viewer (IGV) (108).

Comparisons to Pol II-S5p and H3K27me3 binding in mouse ESCs were performed using GEO data sets GSM2474111 and GSM2474113, respectively.

### RT-qPCR and RNA-seq analysis

Total RNA was extracted and purified from cells directly lysed in TRIzol reagent (Life Technologies). For RT-qPCR, 1 μg of purified RNAs were treated with deoxyribonuclease I (Life Technologies) before subjected to first strand cDNA synthesis using iScript cDNA Synthesis Kit (Bio-Rad) or SuperScript IV VILO Master Mix (Thermo Fisher Scientific) for low abundance transcripts. RT-PCR analysis was carried out with iTaq Universal SYBR Green (Bio-Rad) and gene-specific primers using the CFX Touch Real-Time PCR Detection System (Bio-Rad). Results were normalized to β-actin. Primer sequences are shown in Table S1.

For RNA-seq, total RNA was extracted from cells using TRizol Reagent (Invitrogen). Ribosomal RNAs depletion and libraries preparation were conducted using KAPA HyperPrep kit with Ribo-Erase (Roche) according to manufacturer instructions. Sequencing was performed using Nextseq Illumina sequencer using a NextSeq High-Output 75-cycle kit.

RNA-seq samples were processed using the RNA-seq pipeline implemented in bcbio-nextgen v1.1.3 (https://bcbio-nextgen.readthedocs.org). Raw reads were examined for quality issues using FastQC (98) and aligned the UCSC build mm10 of the mouse genome using STAR (109). Alignments were checked for evenness of coverage, rRNA content, exonic and intronic mapping rates, complexity and other quality metrics using FastQC, Qualimap (110), MultiQC (111) and custom tools. RNA-seq counts were generated using Salmon (112). DESeq2 (113) was used to identify differentially expressed genes with a BH adjusted *P* value cutoff of 0.01; no fold-change cutoff was applied.

### Micrococcal nuclease ChIP-qPCR analysis

MNase-ChIP was performed essentially as described (15) except that ESCs were cross-linked with formaldehyde for 5 min. Cross-linked chromatin were immunoprecipitated with rabbit anti-HA monoclonal antibodies (Cell Signaling Technology). Purified DNA was quantified by real-time PCR with KAPA SYBR FAST qPCR Master Mix (KAPA Biosystems) and gene-specific primers using the CFX Touch Real-Time PCR Detection System (Bio-Rad). Primer sequences are shown in Table S2.

### CRISPR/Cas9-mediated genome editing

For generation of HA-tagged SOX2 and SOX15 knockin ESC lines, single-guide RNA (sgRNA) targeting genomic region immediately upstream of the translation stop codon of *Sox2* or *Sox15* was cloned into LentiCRISPRv2 vector (Addgene). Single-stranded (ss) donor oligonucleotides containing a flexible linker GSGT followed by HA tag sequence and translation stop codon, which is flanked by left and right homology arms of about 60-75 base pairs (bp), were synthesized [Integrated DNA Technologies (IDT)]. Both the LentiCRISPRv2-sgRNA plasmid and the ss donor oligonucleotides were transfected into D3 mouse ESC line using Lipofectamine 3000 (Invitrogen). Transfected cells were selected with puromycin (1.5 μg/ml) for 3 days. Cells were then expanded in the absence of puromycin. Single clones were plated into 96-well plates. Positive clones were identified by PCR and confirmed by sequencing and western blotting. Clones selected for further analysis were confirmed to be puromycin-sensitive and express similar levels of key pluripotency genes such as *Nanog*, *Oct4*, and *Sox2*. See Table S3 for sgRNA and ss donor oligonucleotide sequences.

For generation of *Sox15* knockout ESC lines, sgRNA targeting immediately downstream of the translation start codon or sgRNAs flanking the entire *Sox15* gene locus were cloned into LentiCRISPRv2 vector. See Table S4 for sgRNA sequences.

To delete the −17 kb region upstream of the *Hes5* gene locus, sgRNAs flanking a ~1 kb region containing the SOX15 ChIP-seq peaks were cloned into LentiCRISPRv2 vector. Transfection and selection of knockout clones were as described for SOX2 and SOX15 HA-knockin lines. Homozygous deletion clones were confirmed by genomic PCR and sequencing. See Table S5 and S6 for sgRNA sequences and primer sequences for genomic PCR, respectively.

### Colony formation assay

200 cells of WT and *Sox15* KO clones (KO1 and KO2) were plated on 24-well plates either under medium containing serum and LIF or serum-free 2i/LIF medium. After 6 days, cells were fixed and stained for AP activity (EMD Millipore). AP-positive cells were counted and analyzed.

### Immunofluorescence microscopy

WT and *Sox15* KO clones (KO1 and KO2) ESCs were plated on cover glasses and induced to undergo neural differentiation by changing ESC medium to serum-free N2B27 medium. At day 6, cells were fixed in 4% paraformaldehyde for 10 min at room temperature (RT) and permeabilized with 0.1% triton X-100 in PBS for 10 min at RT. After three washes in PBS for 5 min each, cells were blocked with 5% bovine serum albumin (BSA) and 5% goat serum for 1hr at RT and incubated with primary antibodies (anti-βIII-tubulin antibody [BioLegend]) overnight at 4°C. Following three washes, cells were incubated with secondary antibodies (Goat anti-Rabbit IgG Alexa Fluor-488, Life Technologies, 1:500) for 1 hr at RT. Cover glasses were mounted on slides with mountant with 4’,6-diamidino-2-phenylindole (DAPI) (Life Technologies) and imaged with a confocal microscope.

## Statistical analysis

To determine statistical significance, *P* values were calculated by using unpaired two-sided Student’s *t* test. *P* values less than 0.05 (<0.05) were considered as statistically significant, and they were indicated with * (\**p* < 0.05). All data represent the mean ± SEM (error bars).

## Data Availability

The data that support the findings of this study are available within the manuscript and associated supplementary data files. The ChIP-seq and RNA-seq data have been deposited in NCBI’s Gene Expression Omnibus and can be accessed through GEO series accession number GSE200688. Go to https://www.ncbi.nlm.nih.gov/geo/query/acc.cgi?acc=GSE200688 Enter token ahyvyoimrhilvgx into the box to access data.

## Supporting information

Supporting information can be found online.

## Acknowledgments

We thank G. Dailey for help with expression constructs and S. Zheng for tissue culture assistance; We thank S. Agarwal, Z. Zhang, C. Cattoglio and members of the Fong lab for valuable discussion.

