## Supplementary Figures for "SOX15 regulates stem cell pluripotency and promotes neural fate during differentiation by activating *Hes5*"

### Figure S1

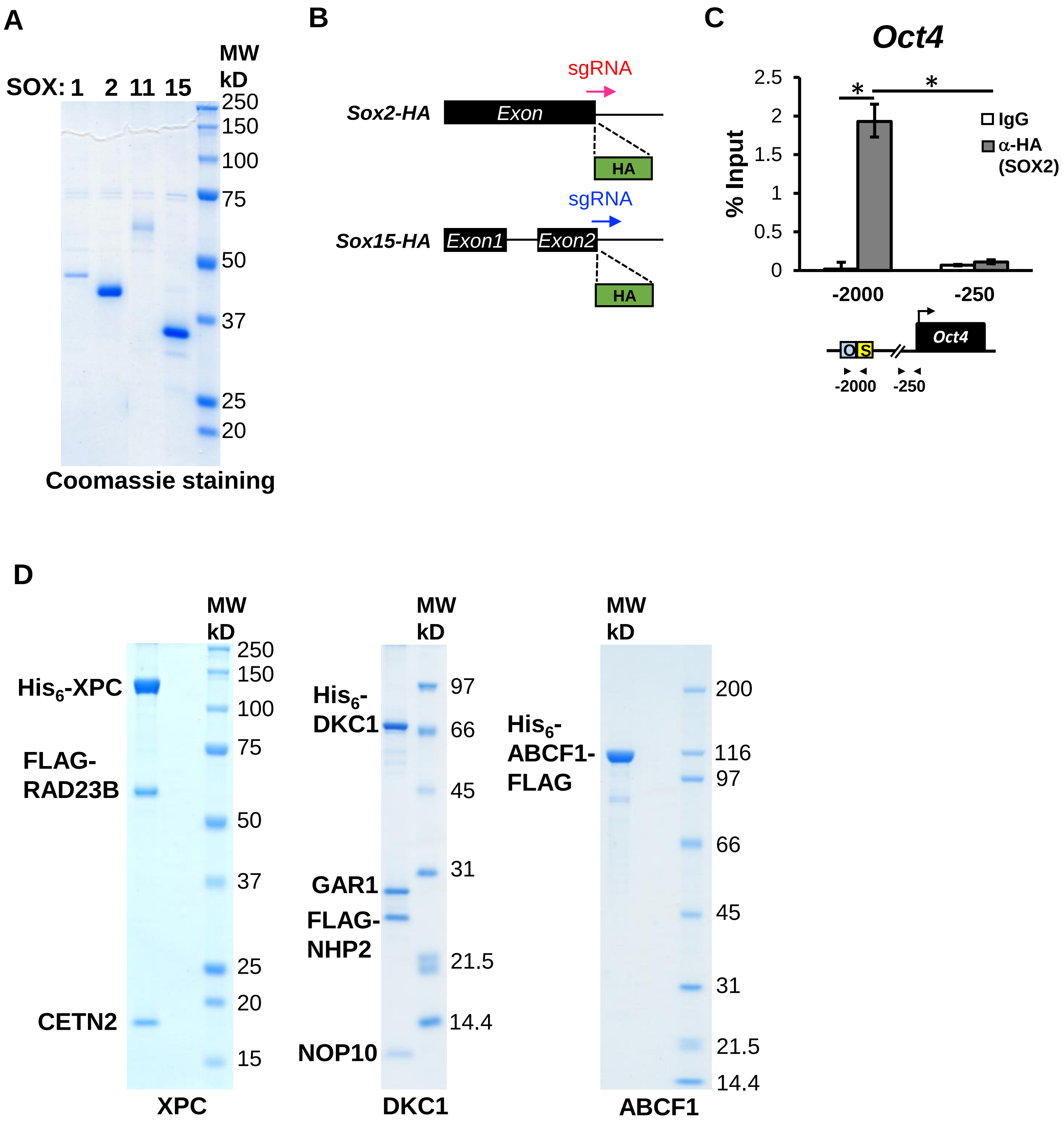
A
SOX: 1 2 11 15
MW kD
250
150
100
75
50
37
25
20
Coomassie staining
B
sgRNA
Sox2-HA
Exon
HA
sgRNA
Sox15-HA
Exon1
Exon2
HA
C
Oct4
% Input
2.5
2
1.5
1
0.5
0
*
*
IgG
α-HA (SOX2)
-2000
-250
O
S
Oct4
-2000
-250
D
MW kD
250
150
100
75
50
37
25
20
15
His6-XPC
FLAG-RAD23B
CETN2
XPC
MW kD
97
66
45
31
21.5
14.4
His6-DKC1
GAR1
FLAG-NHP2
NOP10
DKC1
MW kD
200
116
97
66
45
31
21.5
14.4
His6-ABCF1-FLAG
ABCF1

**Figure S1. Purified SOX proteins and recombinant XPC, DKC1 and ABCF1. (A)** SDS-PAGE and Coomassie staining of FLAG-tagged SOX transcription factors (SOX1, SOX2, SOX11, and SOX15) purified from transiently transfected HeLa cells. **(B)** Schematic diagram depicting CRISPR/Cas9-mediated knockin of an HA-epitope tag to the C-terminus of *Sox2* or *Sox15* gene locus in mouse ESCs. sgRNAs targeting sequences immediately upstream of the translation stop codon are used. **(C)** The anti-HA antibody robustly enriches SOX2-HA at the enhancer but not control region of the *Oct4* locus in SOX2-HA knockin ESCs. MNase-ChIP and data analysis are performed as in Figure 1C. **(D)** SDS-PAGE and Coomassie staining of purified recombinant XPC and DKC1 complexes from insect Sf9 cells, and recombinant ABCF1 from *E. coli*. Subunits of the XPC and DKC1 complexes and affinity-tags used for protein purification are indicated.

### Figure S2

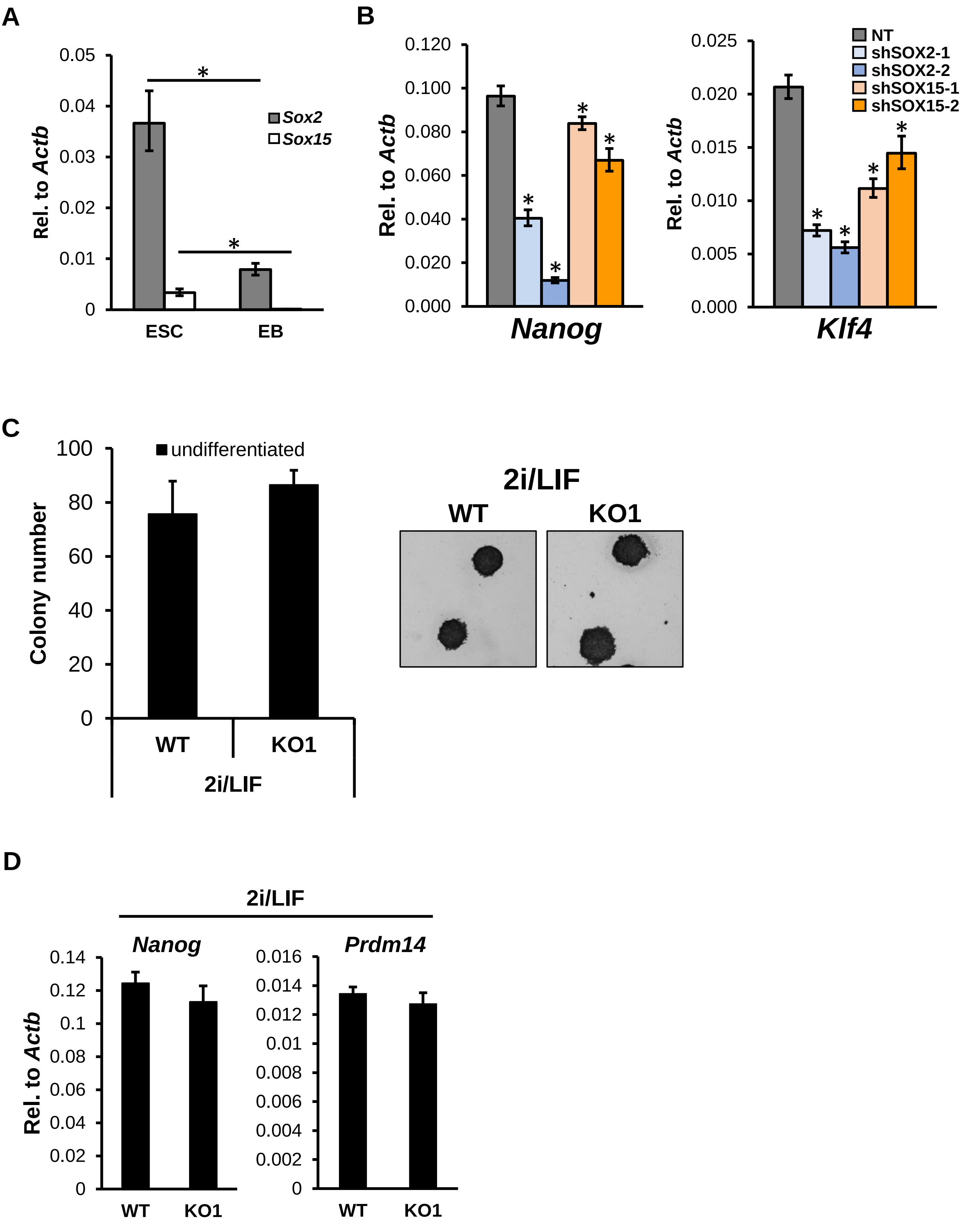
A
B
C
D
Rel. to Actb
0.05
0.04
0.03
0.02
0.01
0
*
Sox2
Sox15
ESC
EB
0.120
0.100
0.080
0.060
0.040
0.020
0.000
Nanog
0.025
0.020
0.015
0.010
0.005
0.000
Klf4
NT
shSOX2-1
shSOX2-2
shSOX15-1
shSOX15-2
Colony number
100
80
60
40
20
undifferentiated
WT
KO1
2i/LIF
0.14
0.12
0.1
0.08
0.06
0.04
0.02
Prdm14
0.016
0.014
0.012
0.01
0.008
0.006
0.004
0.002

**Figure S2. SOX15 depletion compromises pluripotency gene expression in ESCs grown in serum/LIF condition. (A)** *Sox2* and *Sox15* mRNA levels are downregulated in differentiating embryoid bodies (EB) compared to ESCs, determined by qPCR and analyzed as in Figure 2A. **(B)** qPCR analyses of *Nanog* and *Klf4* mRNA levels in SOX2 and SOX15-knockdown ESCs. **(C)** Self-renewal defect in *Sox15* KO ESCs upon single cell dissociation is rescued by culturing dissociated ESCs in 2i/LIF condition. Representative images of colony morphology are shown (right). **(D)** qPCR analyses of *Nanog* and *Prdm14* mRNA levels in WT and *Sox15* KO (KO1) cultured in 2i/LIF condition, relative to *Actb*. Error bars represent SEM. $n$ = 3. (*) $P < 0.05$, calculated by two-sided Student's t-test.

### Figure S3

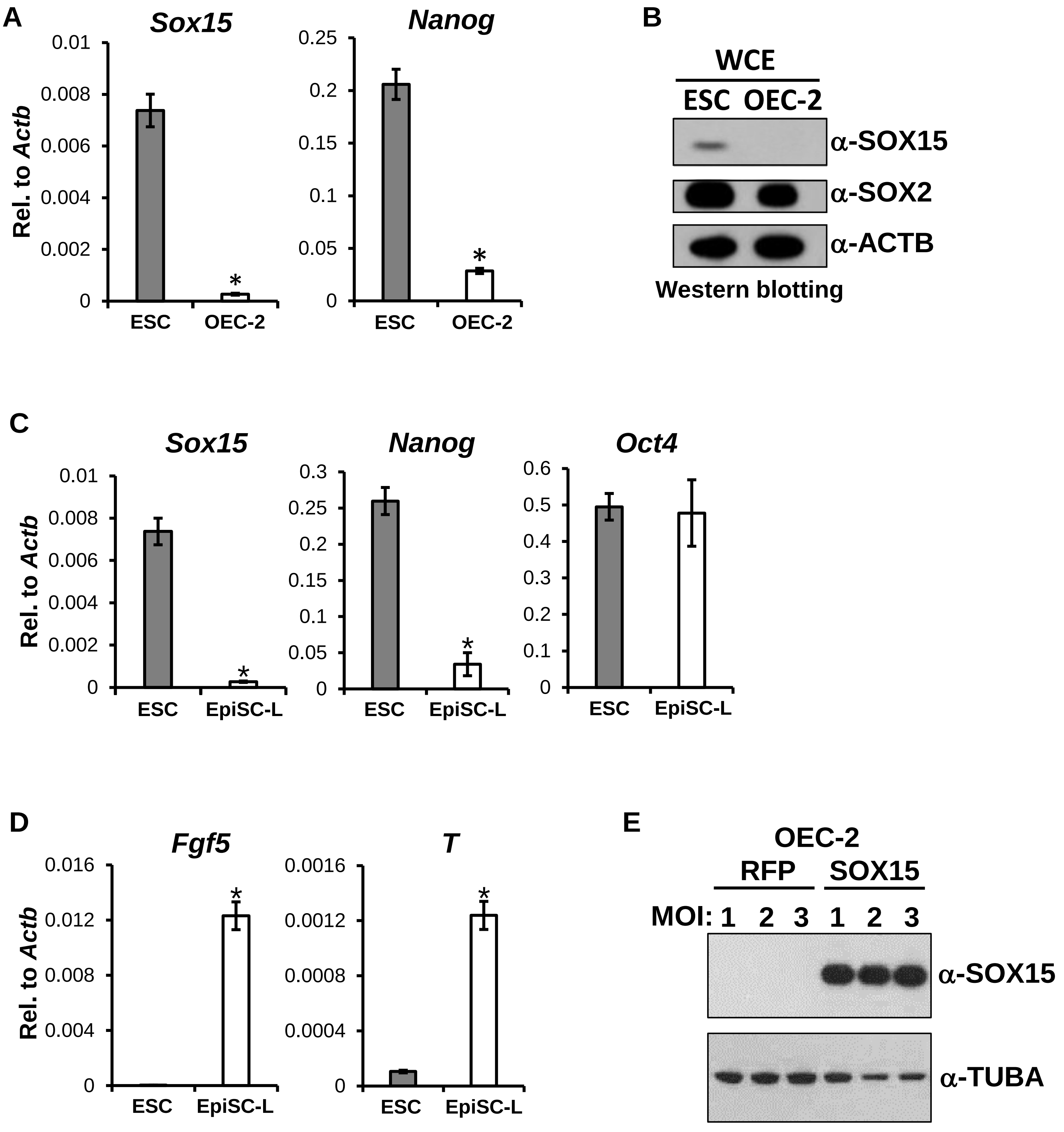
A
Sox15
Nanog
Rel. to Actb
ESC
OEC-2
B
WCE
ESC OEC-2
α-SOX15
α-SOX2
α-ACTB
Western blotting
C
Sox15
Nanog
Oct4
Rel. to Actb
ESC
EpiSC-L
D
Fgf5
T
Rel. to Actb
ESC
EpiSC-L
E
OEC-2
RFP
SOX15
MOI: 1 2 3 1 2 3
α-SOX15
α-TUBA

**Figure S3. SOX15 expression levels in naive and primed pluripotent states. (A)** *Sox15* and *Nanog* mRNA levels in ESCs cultured in 2i/LIF and EpiSC line OEC-2 are analyzed by qPCR, normalized to *Actb*. **(B)** Western blotting of WCEs from ESCs and OEC-2 cells using antibodies against SOX15 and SOX2; ACTB as loading control. **(C)** Pluripotency-associated genes *Nanog*, *Sox15*, and *Oct4* mRNA levels in ESCs and EpiSC-like cells (EpiSC-L) converted from ESCs in vitro are analyzed by qPCR, normalized to *Actb*. **(D)** EpiSC-enriched genes (*Fgf5* and *T*) are upregulated in EpiSC-L compared to ESCs, determined by qPCR and normalized to *Actb*. **(E)** Expression level of exogenous SOX15 in OEC-2 cells transduced with lentiviruses expressing RFP or SOX15 at indicated MOI is shown by western blotting of WCEs using antibodies against SOX15 and TUBA as loading control. Error bars represent SEM. $n = 3$. (*) $P < 0.05$, calculated by two-sided Student's t-test.

### Figure S4

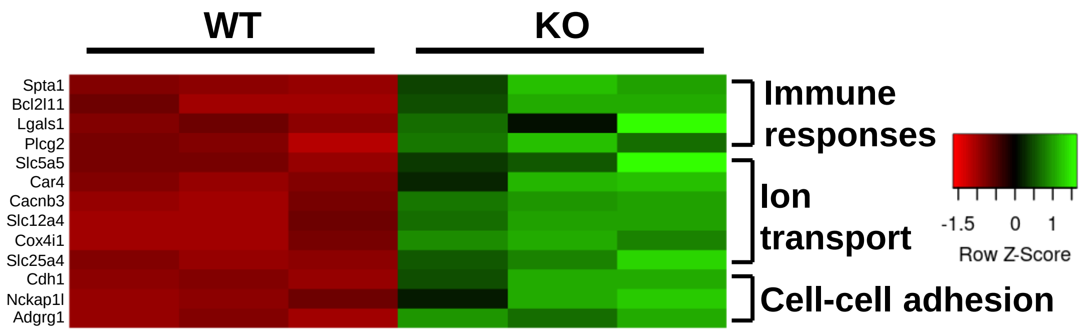


**Figure S4. Upregulated genes in *Sox15*-knockout ESCs.** Heatmap showing the relative expression levels of genes belonging to the indicated categories in three biological replicates each of WT and *Sox15* KO1 ESCs cultured in serum/LIF. Scaled values indicate relative downregulation (red) or upregulation (green) of gene expression.

### Figure S5

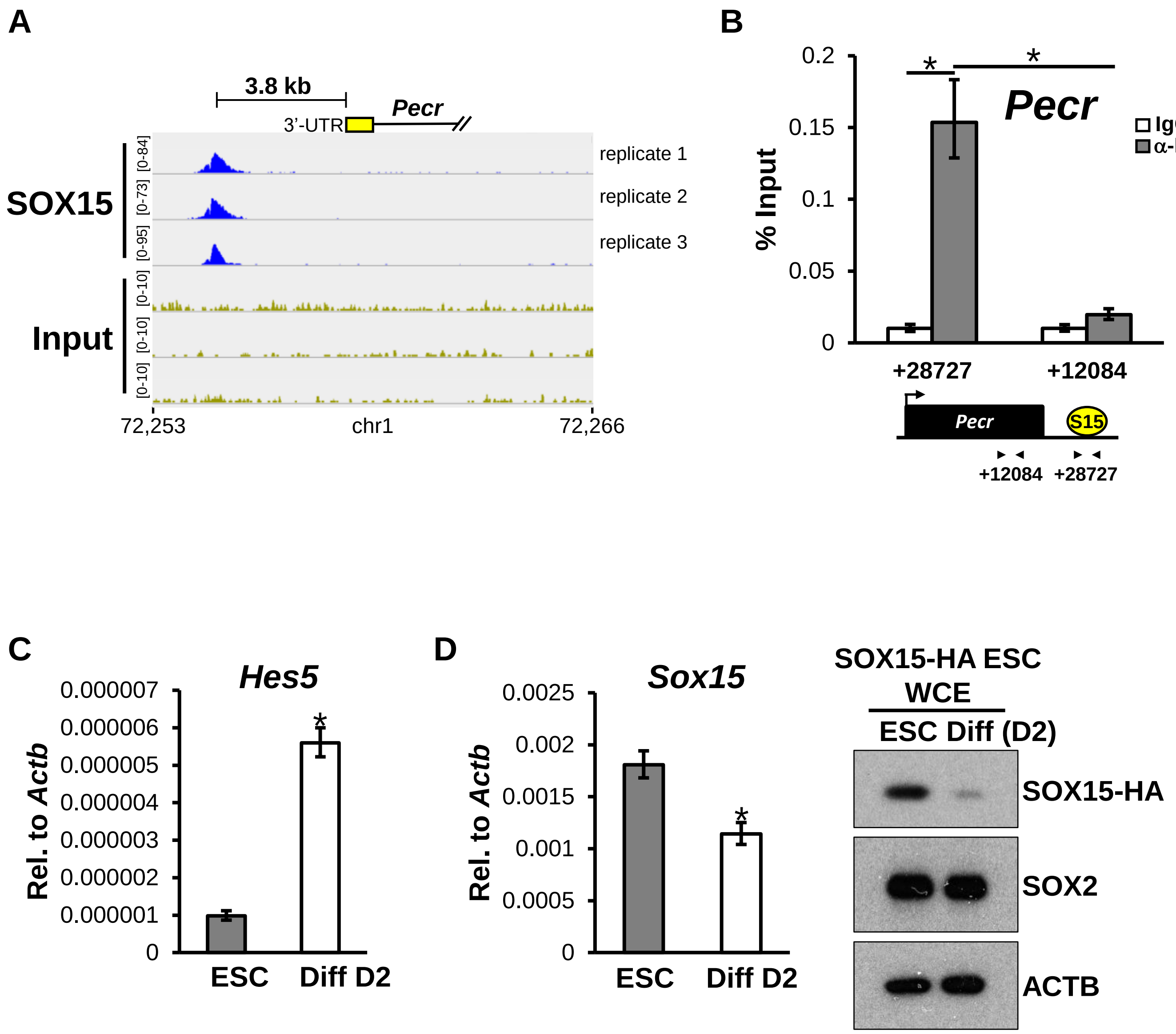
A
3.8 kb
Pecr
3'-UTR
SOX15
Input
[0-84]
[0-73]
[0-95]
[0-10]
[0-10]
[0-10]
replicate 1
replicate 2
replicate 3
72,253
chr1
72,266
B
Pecr
% Input
0.2
0.15
0.1
0.05
0
IgG
α-HA
+28727
+12084
S15
C
Hes5
Rel. to Actb
0.000007
0.000006
0.000005
0.000004
0.000003
0.000002
0.000001
0
ESC
Diff D2
D
Sox15
0.0025
0.002
0.0015
0.001
0.0005
SOX15-HA ESC
WCE
ESC Diff (D2)
SOX15-HA
SOX2
ACTB

**Figure S5. SOX15 occupancy in ESCs and expression level upon neural differentiation. (A)** IGV-computed SOX15-HA ChIP-seq tracks from three biological replicates are plotted for the *Pecr* gene locus as described in Figure 5A. One of the strongest SOX15 ChIP-seq peaks is detected at ~3.8 kb downstream of the 3' end of *Pecr*. **(B)** Validation of SOX15-HA ChIP-seq results. MNase-ChIP analysis of SOX15 occupancy on control and bound regions near the *Pecr* gene in mouse ESCs. Relative positions of SOX15-bound (S15) and control regions relative to TSS are shown. Enrichment and data analysis are as described in Figure 1C. **(C)** qPCR analysis showing upregulation of *Hes5* mRNA in SOX15-HA knockin ESCs after 2 days of differentiation (Diff D2). *Hes5* transcript levels are normalized to *Actb*. **(D)** *Sox15* mRNA levels are downregulated at day 2 of neural differentiation, determined by qPCR and normalized to *Actb* (left). Western blotting of WCEs of undifferentiated (ESC) and differentiating (Diff D2) ESCs, using antibodies against HA (SOX15-HA), SOX2, and ACTB (right). SOX2 level remains largely unchanged. Error bars represent SEM. $n = 3$. (*) $P < 0.05$, calculated by two-sided Student's t-test.

#### Figure S6

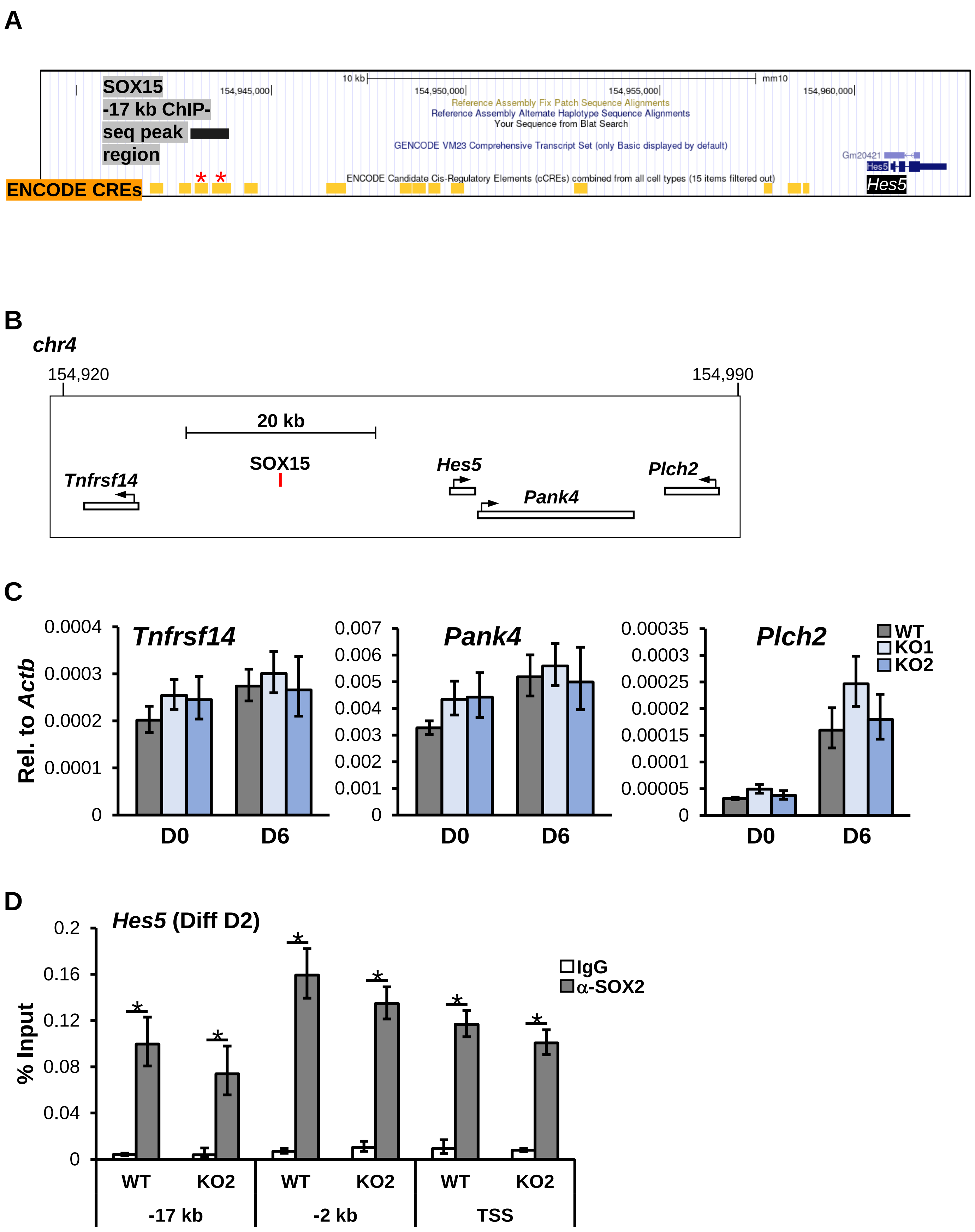
A
SOX15 -17 kb ChIP-seq peak region
ENCODE CREs
10 kb
mm10
154,945,000
154,950,000
154,955,000
154,960,000
Reference Assembly Fix Patch Sequence Alignments
Reference Assembly Alternate Haplotype Sequence Alignments
Your Sequence from Blat Search
GENCODE VM23 Comprehensive Transcript Set (only Basic displayed by default)
ENCODE Candidate Cis-Regulatory Elements (cCREs) combined from all cell types (15 items filtered out)
Gm20421
Hes5
Hes5
B
chr4
154,920
154,990
20 kb
Tnfrsf14
SOX15
Hes5
Pank4
Plch2
C
Rel. to Actb
Tnfrsf14
Pank4
Plch2
WT
KO1
KO2
D0
D6
D
Hes5 (Diff D2)
% Input
IgG
α-SOX2
WT
KO2
-17 kb
-2 kb
TSS

### Figure S6 cont'd

E

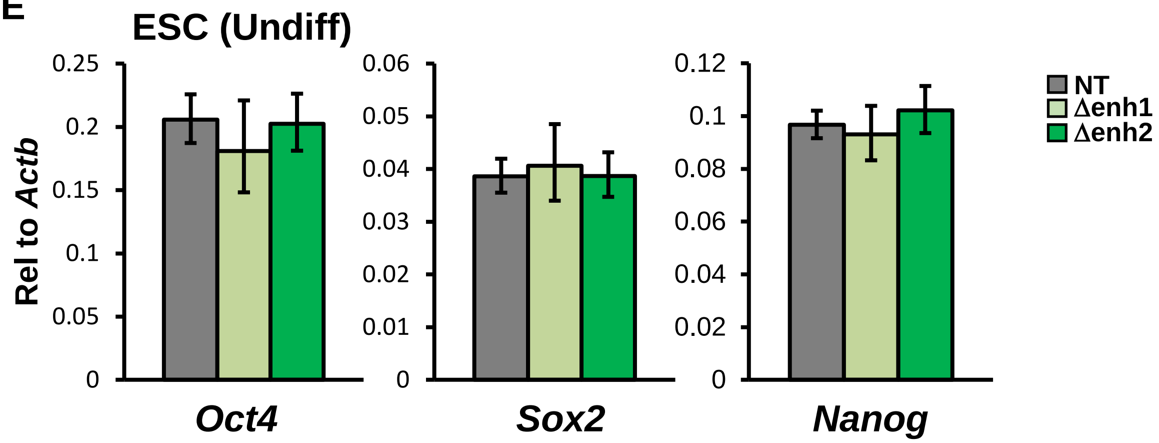


F

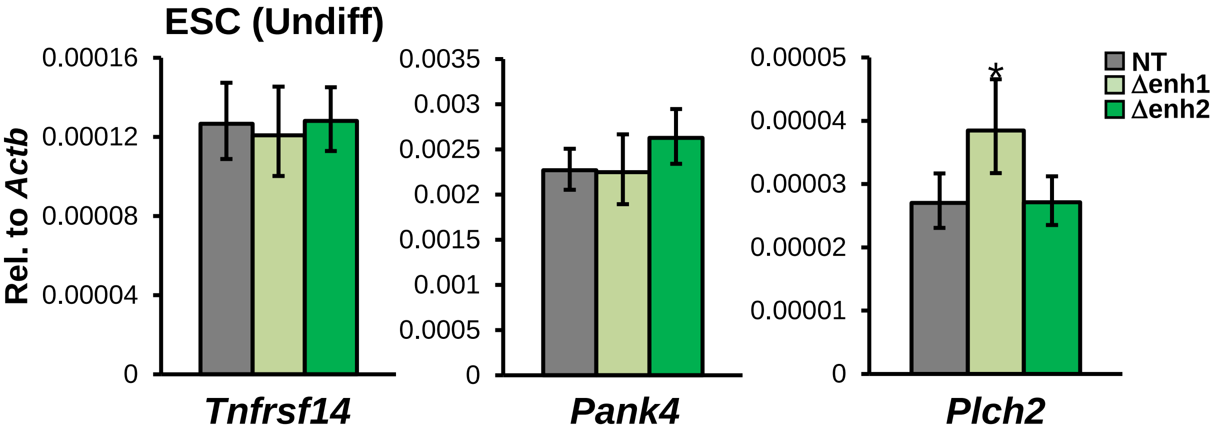


G

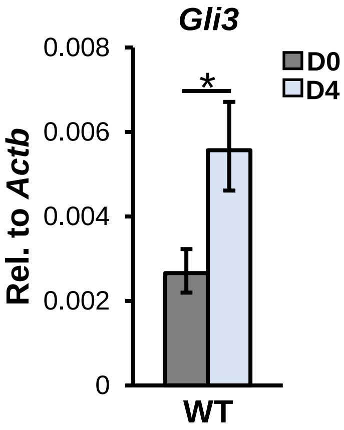


H

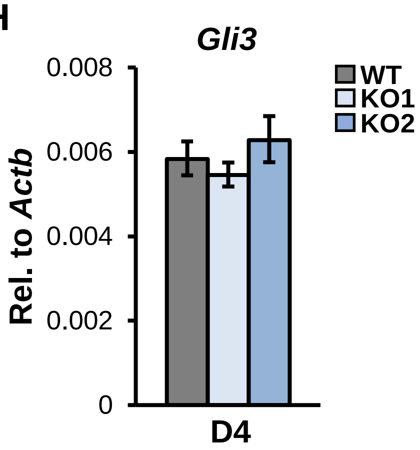

**Figure S6. Deletion of the -17 kb region compromises *Hes5* expression but not genes in its neighborhood. (A)** UCSC Genome Browser generated diagram showing ENCODE cis-regulatory elements (CREs, orange bars) around the -17 kb SOX15 ChIP-seq peak region (black bar) upstream of the *Hes5* gene locus on chromosome (chr) 4. Red asterisks indicate CREs that overlap with SOX15 ChIP-seq peak. These highlighted CREs display distal enhancer-like signatures such as DNase I hypersensitivity and enrichment of H3K27ac and H3K4me3 signals in neural cell types and tissues. **(B)** Position of genes near the -17 kb region (*Tnfrsf14*, *Hes5*, *Pank4*, and *Plch2*) within a ~70 kb window in chr 4 is shown. SOX15-bound region -17 kb upstream of *Hes5* is marked in red. Genomic coordinates are in kilobases. **(C)** Expression of *Tnfrsf14*, *Pank4*, and *Plch2* in undifferentiated ESCs (D0) and day 6 differentiated ESCs (D6) is not affected by SOX15 depletion in KO ESCs. mRNA levels in WT and *Sox15* KO (KO1 and 2) ESCs are analyzed by qPCR, normalized to *Actb*. **(D)** Enrichment of SOX2 at the *Hes5* locus is not dependent on SOX15. Occupancy of SOX2 at -17 kb, -2 kb, and TSS of the *Hes5* gene in WT and *Sox15* KO ESCs (KO2) at day 2 of differentiation (Diff D2) is analyzed by MNase-ChIP, using antibodies against SOX2 and Ig as control. **(E)** Expression of core pluripotency genes *Oct4*, *Sox2*, *and Nanog* is not affected by deletion of the -17 kb region. Their respective mRNA levels in WT and ∆enh ESC lines are quantified by qPCR, normalized to *Actb*. **(F)** Deletion of the -17 kb region does not compromise *Tnfrsf14*, *Pank4*, and *Plch2* expression in ESCs. Their mRNA levels in undifferentiated ESCs are analyzed as in (D). **(G)** *Gli3* mRNA levels in undifferentiated (D0) and day 4 differentiated (D4) WT ESCs are analyzed by qPCR, normalized to *Actb*. **(H)** qPCR analysis of *Gli3* mRNA levels in WT, *Sox15* KO1 and KO2 ESCs at D4 of differentiation, normalized to *Actb*. Error bars represent SEM. $n = 3$. (*) $P < 0.05$, calculated by two-sided Student's t-test.
