## Supplementary Table for "SOX15 regulates stem cell pluripotency and promotes neural fate during differentiation by activating *Hes5*"

**Supplementary Tables**

**Table S1. Primers used for RT-qPCR analysis**

| **Gene** | **Forward** | **Reverse** |
| --- | --- | --- |
| *Nanog* | CCTCAGCCTCCAGCAGATGC | CCGCTTGCACTTCATCCTTTG |
| *Klf4* | CAGTGGTAAGGTTTCTCGCC | GCCACCCACACTTGTGACTA |
| *Prdm14* | GCATCCTGGTTCCCACAGAG | GCTGCAGAACACGCCAAAGTG |
| *Esrrb* | TACCTGAACCTGCCGATTTC | CCCAGTTGATGAGGAACACA |
| *Nr5a2* | GTTGAGTGGGCCAGGAGTAG | GATTGTCCCTTCCTTCCCAT |
| *Pou5f1(Oct4)* | GAAGCAGAAGAGGATCACCTTG | TTCTTAAGGCTGAGCTGCAAG |
| *Sox2* | CCGGCACGGCCATTAAC | CCCTCCCAATTCCCTTGTAT |
| *Sox15* | CAACTACCCAAGGGAGCAGA | GAGGGAAGGTGCATGGTAAG |
| *Fgf5* | TTGCGACCCAGGAGCTTAAT | CTACGCCTCTTTATTGCAGCAT |
| *Nes* | TCCCTTAGTCTGGAAGTGGCTA | GGTGTCTGCAAGCGAGAGTT |
| *Sox1* | GTGACATCTGCCCCCATC | GAGGCCAGTCTGGTGTCAG |
| *Hes1* | CGGCATTCCAAGCTAGAGAAGG | GGTAGGTCATGGCGTTGATCTG |
| *Hes5* | CTTCTGCGAAGTTCCTGGTC | ATGTGGACCTTGAGGTGAGG |
| *T* | CTCTAATGTCCTCCCTTGTTGCC | TGCAGATTGTCTTTGGCTACTTTG |
| *Mash1* | TCTCCTGGGAATGGACTTTG | CGTTGGCGAGAAACACTAAAG |
| *Pax6* | GTTCCCTGTCCTGTGGACTC | ACCGCCCTTGGTTAAAGTCT |
| *Tnfrsf14* | ATGCAAATGGCCTGAGCAAG | ATCTGCACACAGTGTCCTTCC |
| *Pank4* | TGGAGCCTACAAGTTCAAGGAC | GCACCCTTTAATCAAGCAGGTC |
| *Plch2* | TGATGAGATGGAGGATGACTGC | CCTTGATCAGAGAATCCAGCTTC |
| *Actb* | AGAGGGAAATCGTGCGTGAC | CAATAGTGATGACCTGGCCGT |

**Table S2. Primers used for MNase ChIP-qPCR analysis**

| **Name** | **Forward** | **Reverse** |
| --- | --- | --- |
| *Oct4-250* | AGCAACTGGTTTGTGAGGTGTCCGGTGAC | TCCCCAATCCCACCCTCTAGCCTTGAC |
| *Oct4-2000* | GGAACTGGGTGTGGGGAGGTTGTA | AGCAGATTAAGGAAGGGCTAGGACGAGAG |
| *Sox2-3581* | CCCTGTTCCAAGTCTCTTTCTG | GATTTCAATCCAACACCATCATAG |
| *Sox2+3513* | TTTTCGTTTTTAGGGTAAGGTACTGG | CGTGAATAATCCTATATGCATCACAAT |
| *Nanog-952* | GGCAAACTTTGAACTTGGGATGTGGAAATA | CTCAGCCGTCTAAGCAATGGAAGAAGAAAT |
| *Nanog-199* | TCTGGGTCACCTTACAGCTTC | TCACAGTTAATCCCACCTGCAG |
| *Pecr+12084* | CTCTGACCCAACCCCAGTAA | CACTCGCACAGAGATGGAGA |
| *Pecr+28727* | GGAACCTATGCCTCCTCTCC | CGTGGCGAGTGGTATTCTTT |
| *Hes5 TSS* | AACTGGCAATAAAGCGCCTA | CATTCGCAGGTACAGGATGA |
| *Hes5-2 kb* | CTGCCCCCTCAACTACTGTC | CATAGGGCAGGATTGGAGTC |
| *Hes5 -17 kb* | AATCCTAGGGCACAATGGACTG | AGAGACTTTGAGGCTGCCTAC |

**Table S3. sgRNA and ss donor oligo sequences for generating endogenously HA-tagged *Sox2* and *Sox15* knock-in mouse ESC lines**

| **Name** | **Sequence** |
| --- | --- |
| ***Sox2*:** |  |
| sgRNA-F | CACCGGTGCCGTTAATGGCCGTGCC |
| sgRNA-R | AAAGGCACGGCCATTAACGGCACC |
| ss donor oligo | GAGCCCGCTGCGCCCAGTAGACTGCACATGGCCCAGCACTACCAGAGCGGCCCGGTGCCCGGCACGGCCATTAACGGCACACTGCCCCTGTCGCACATGGGATCCGGTACCTACCCATACGATGTTCCAGATTACGCTTGAGGGCTGGACTGCGAACTGGAGAAGGGGAGAGATTTTCAAAGAGATACAAGGGAATTGGG |
| ***Sox15*:** |  |
| sgRNA-F | CACCGTGGCATGGGGGCTCCAGCAA |
| sgRNA-R | AAATTGCTGGAGCCCCCATGCCAC |
| ss donor oligo | AGCTAAGACCCTCTTTCTCCCCTTACCTATCCCCAGACTCTTCCACTCCATATAATACTTCCCTTGCTGGAGCCCCCATGCCAGTAACCCACCTTGGATCCGGTACCTACCCATACGATGTTCCAGATTACGCTTAACTGGCAGGGACGCCCAGGGCCAACCCCACAGACCCCTACAGATCTCCAAGGAACCAGAATGCA |

**Table S4. sgRNA sequences for generating *Sox15* knockout in mouse ESCs**

| **Name** | **Sequence** |
| --- | --- |
| sgRNA-1F | CACCGTCTGGAGCCCTTTACCGGAG |
| sgRNA-1R | AAACCTCCGGTAAAGGGCTCCAGAC |
| sgRNA-2F | CACCGCGGGAGCCCTGGAGCGTCTG |
| sgRNA-2R | AAACAGACGCTCCAGGGCTCCCGC |
| sgRNA-3F | CACCGTGGCATGGGGGCTCCAGCAA |
| sgRNA-3R | AAATTGCTGGAGCCCCCATGCCAC |

**Table S5. sgRNA sequences for generating *Hes5* -17 kb deletion mouse ESC lines**

| **Name** | **Sequence** |
| --- | --- |
| sgRNA-1F | CACCGGCGGTGCATTTAACAAGTAAGGG |
| sgRNA-1R | AAACCCCTTACTTGTTAAATGCACCGCC |
| sgRNA-2F | CACCGGACGTGAAGTGAAAGTACCCAGG |
| sgRNA-2R | AAACCCTGGGTACTTTCACTTCACGTCC |

**Table S6. Genomic PCR primer sequences for screening *Hes5* -17 kb deletion mouse ESC lines**

| **Name** | **Sequence** |
| --- | --- |
| Hes5-17-F | CCATCGAGTCACCACAGTAAAA |
| Hes5-17-R | ACCTTGAGCAGTCTCTGACTATCTT |
